## Supplementary materials for "Estimating the proportion of beneficial mutations that are not adaptive in mammals"

### 1 Mutation-selection codon models

#### 1.1 Prior distributions and parameters of the model

The parameterization of the models is described as a Bayesian hierarchical model, including the prior distributions and the parameters of the model. This hierarchical model is formally represented as directed acyclic graph of dependencies between variables, depicted below.

Figure S1: Bayesian hierarchical model.

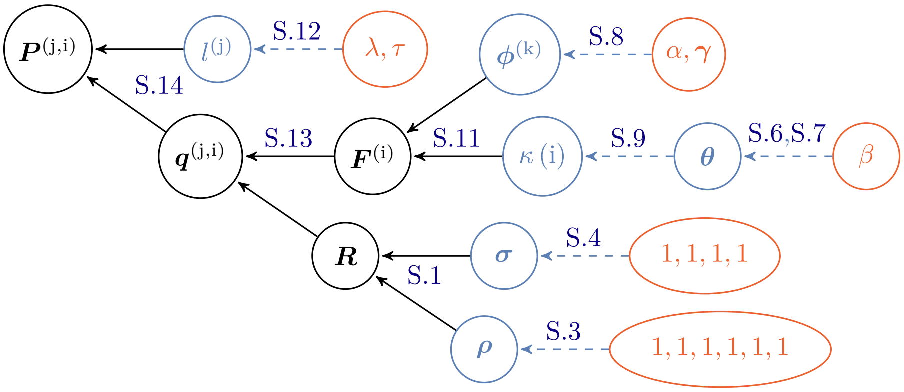

Nodes of the directed acyclic graph are the variables, and edges are the functions. Hyper-parameters are depicted in red circles, random variables in blue circles, and transformed variables in black. Blue dashed line denotes a drawing from a random distribution, and black solid lines denote a function. All the nodes pointing toward a given node (upstream) are its dependencies which determine its distribution. The other way around, following the arrows in the DAG (downstream), simple prior distributions are combined together to form more complex joint prior distributions which ultimately define the prior distribution of the model.

#### 1.2 Nucleotide mutation rates

The generalized time-reversible (GTR) nucleotide mutation rate matrix $\boldsymbol{R}$ is a function of the nucleotide frequencies $\boldsymbol{\sigma}$ and the symmetric exchangeability rates $\boldsymbol{\rho}$ [1]. $\boldsymbol{\sigma} = (\sigma_A, \sigma_C, \sigma_G, \sigma_T)$ is the equilibrium base frequency vector, giving the frequency at which each base occurs at each site. $\boldsymbol{\rho} = (\rho_{AC}, \rho_{AG}, \rho_{AT}, \rho_{CG}, \rho_{CT}, \rho_{GT})$ is the vector of exchangeabilities between nucleotides. Altogether, the rate matrix is:

$$\boldsymbol{R} = \begin{array}{c} \\ A \\ C \\ G \\ T \end{array} \begin{array}{c} \begin{array}{cccc} A & C & G & T \end{array} \\ \begin{pmatrix} -\mu_A & \rho_{AC}\sigma_C & \rho_{AG}\sigma_G & \rho_{AT}\sigma_T \\ \rho_{AC}\sigma_A & -\mu_C & \rho_{CG}\sigma_G & \rho_{CT}\sigma_T \\ \rho_{AG}\sigma_A & \rho_{CG}\sigma_C & -\mu_G & \rho_{GT}\sigma_T \\ \rho_{AT}\sigma_A & \rho_{CT}\sigma_C & \rho_{GT}\sigma_G & -\mu_T \end{pmatrix} \end{array} \tag{S.1}$$

By definition, the sum of the entries in each row of the nucleotide rate matrix $\boldsymbol{R}$ is equal to 0, giving the diagonal entries:

$$\mu_a = \sum_{b \neq a, b \in \{A,C,G,T\}} R_{a,b} \tag{S.2}$$

The prior on the exchangeabilities $\boldsymbol{\rho}$ is a uniform Dirichlet distribution of dimension 6:

$$\boldsymbol{\rho} \sim \text{Dir}\left(1, 1, 1, 1, 1, 1\right). \tag{S.3}$$

The prior on the equilibrium base frequencies $\boldsymbol{\sigma}$ is a uniform Dirichlet distribution of dimension 4:

$$\boldsymbol{\sigma} \sim \text{Dir}\left(1,1,1,1\right) \tag{S.4}$$

The general time-reversible nucleotide matrix is normalized with a total flow of 1:

$$\sum_{a \in \{A,C,G,T\}} -\sigma_a R_{a,a} = 1, \tag{S.5}$$

such that we expect 1 substitution per unit of branch length.

#### 1.3 Site-specific amino-acid fitness profiles

Site-specific amino-acid fitness profiles are assumed i.i.d. from a mixture model, itself endowed with a truncated Dirichlet process prior. Specifically, the mixture has K components (K = 30 by default). The prior on component weights ($\boldsymbol{\theta}$) is modeled using a stick-breaking process, truncated at K and of parameter $\beta = 1$:

$$\begin{gathered} \boldsymbol{\theta} \sim \text{StickBreaking}\left(\text{K}, \beta\right) \\ \iff \theta_{\text{k}} = \psi_{\text{k}} \cdot \prod_{l=1}^{\text{k}-1} \left(1 - \psi_l\right), \ \text{k} \in \{1, \ldots, \text{K}\}, \end{gathered} \tag{S.6}$$

where $\psi_{\text{k}}$ are i.i.d. from a beta distribution

$$\psi_{\text{k}} \sim \text{Beta}\left(1, \beta\right), \ \text{k} \in \{1, \ldots, \text{K}\}. \tag{S.7}$$

Of note, the weights decrease geometrically in expectation, at rate $\beta$, such that lower values of $\beta$ induce more heterogeneous distributions of weights.

Each component of the mixture defines a 20-dimensional fitness profile $\boldsymbol{\phi}^{(\text{k})}$ (summing to 1), for $\text{k} \in \{1, \ldots, \text{K}\}$. These fitness profiles are i.i.d. from a Dirichlet of center $\boldsymbol{\gamma} = 1$ and concentration $\alpha = 1$:

$$\boldsymbol{\phi}^{(\text{k})} \sim \text{Dir}\left(\boldsymbol{\gamma},\ \alpha\right), \ \text{k} \in \{1, \ldots, \text{K}\}. \tag{S.8}$$

Site allocations to the mixture components $\kappa\left(\text{i}\right) \in \{1, \ldots, \text{K}\}$, for $\text{i} \in \{1, \ldots, \text{N}\}$ running over the N sites of the alignment, are i.i.d. multinomial of parameter $\boldsymbol{\theta}$:

$$\boldsymbol{m} \sim \text{Multinomial}\left(\boldsymbol{\theta}\right), \tag{S.9}$$

$$\text{where } m_{\text{k}} = \sum_{\text{i} \in \{1, \ldots, \text{N}\}} \mathbb{1}_{\kappa(\text{i})=\text{k}} \tag{S.10}$$

For a given parameter configuration for the mixture, the scaled fitness $\boldsymbol{F}^{(\text{i})}$ at site i, are obtained by taking the logarithm of fitness assigned to this site:

$$\boldsymbol{F}^{(\text{i})} = \ln\left(\boldsymbol{\phi}^{(\kappa(\text{i}))}\right), \ \text{i} \in \{1, \ldots, \text{N}\}. \tag{S.11}$$

#### 1.4 Branch length

The topology of the rooted phylogenetic tree is supposed to be known and is not estimated by the model. The branch lengths $l^{(\text{j})}$ are defined as the expected number of neutral substitutions per DNA site along a branch, each from a Gamma distribution of mean $\lambda = 0.1$ and scale $\tau = 1$:

$$l^{(\text{j})} \sim \text{Gamma}\left(\lambda, \tau\right). \tag{S.12}$$

#### 1.5 Codon substitution rates

The mutation rate between codons $a$ and $b$, denoted $\mu_{a \mapsto b}$ depends on the underlying nucleotide change between the codons. First, if codons $a$ and $b$ are not nearest-neighbours, $\mu_{a \mapsto b}$ is equal to 0. Second, if codons $a$ and $b$ are only one mutation away, $\mu_{a \mapsto b}$ is given by the underlying nucleotide relative rate ($R_{a \mapsto b}$).

For a given site i, the codon substitution rate matrix $\boldsymbol{q}^{(\mathrm{i})}$ is given by:

$$\begin{cases} q^{(\mathrm{i})}_{a \mapsto b} = 0 \text{ if codons } a \text{ and } b \text{ are not nearest-neighbors,} \\ q^{(\mathrm{i})}_{a \mapsto b} = \mu_{a \mapsto b} \text{ if codons } a \text{ and } b \text{ are synonymous,} \\ q^{(\mathrm{i})}_{a \mapsto b} = \mu_{a \mapsto b} \dfrac{F^{(\mathrm{i})}_b - F^{(\mathrm{i})}_a}{1 - \mathrm{e}^{F^{(\mathrm{i})}_a - F^{(\mathrm{i})}_b}} \text{ if } a \text{ and } b \text{ are non-synonymous,} \\ q^{(\mathrm{i})}_{a,a} = - \displaystyle\sum_{b \neq a, b=1}^{61} q^{(\mathrm{i})}_{a \mapsto b}. \end{cases} \tag{S.13}$$

Together, the probability of transition between codons for a given branch j and site i is:

$$\boldsymbol{P}^{(\mathrm{j},\mathrm{i})} = \mathrm{e}^{l^{(\mathrm{j})} \boldsymbol{q}^{(\mathrm{i})}}, \tag{S.14}$$

which are the matrices necessary to compute the likelihood of the data ($D$) given the parameters of the model using the pruning algorithm.

#### 1.6 Bayesian implementation

Bayesian inference was conducted using Markov Chain Monte Carlo (MCMC). Most phylogenetic MCMC samplers target the distribution over the model parameters given the sequence alignment, which means that they have to repeatedly invoke the pruning algorithm to recalculate the likelihood which is most often the limiting step of the MCMC. An alternative, which is used here, is to do the MCMC conditionally on the detailed substitution history $\mathcal{H}$, thus doing the MCMC over the augmented configuration ($\mathcal{H}$, $D$), under the target distribution obtained by combining the mapping-based likelihood with the prior over model parameters. The key idea that makes this strategy efficient is that the mapping-based likelihood depends on compact summary statistics of $\mathcal{H}$, leading to very fast evaluation of the likelihood. On the other hand, this requires to implement more complex MCMC procedures that have to alternate between:

1. sampling $\mathcal{H}$ conditionally on the data and the current parameter configuration.
2. re-sampling the parameters conditionally on $\mathcal{H}$.

To implement the mapping-based MCMC sampling strategy, we first sample the detailed substitution history $\mathcal{H}$ for all sites along the tree. Several methods exist for doing this [2, 3], which are used here in combination (first trying the accept-reject method of Nielsen [2], then switching to the uniformization approach of Rodrigue *et al.* [3] if the first round has failed).

Then, we write down the probability of $\mathcal{H}$ given the parameters, and finally, we collect all factors that depend on some parameter of interest and make some simplifications. This ultimately leads to relatively compact sufficient statistics allowing for fast numerical evaluation of the likelihood [4, 5].

##### 1.7 BayesCode software

In *BayesCode* (github.com/ThibaultLatrille/BayesCode, v1.3.1), we ran the mutation-selection codon models *mutselomega* for 2000 points of MCMC with the options:

```
mutselomega ---omegashift 0.0 --ncat 30 -a my_alignment.phy -t my_tree.newick -u 2000 my_genename
```

The collection of site-specific fitness profiles ($\boldsymbol{F}^{(i)}, \forall i$) are then obtained by running *readmutselomega*, reading 1000 points of MCMC (first 1000 are considered as burn-in) with the options:

```
readmutselomega --every 1 --until 2000 --burnin 1000 --ss my_genename
```

The gene-specific $4 \times 4$ nucleotide mutation rate matrix ($\boldsymbol{\mu}$) is also obtained by running *readmutselomega*, reading 1000 points of MCMC (first 1000 are considered as burn-in) with the options:

```
readmutselomega --every 1 --until 2000 --burnin 1000 --nuc my_genename
```

### 2 Beneficial mutations in the terminal lineages and populations

#### 2.1 Example sites

We extracted protein-coding DNA alignments across mammals centered around the codon site (two flanking codons in white background) for which the beneficial non-adaptive mutations ($\mathcal{B}_0$) have been detected in *Chlorocebus sabaeus*. We also show the translated amino acids of this region. For instance, in DNA alignment of gene SELE shown below, the nucleotide at site 1722 has mutated (from T to C) at the basis of Simiiformes (monkeys and apes), modifying the corresponding amino acid from Serine to Proline, but has been subsequently reverted in the branch of *Chlorocebus sabaeus*. However, other substitutions classified as $\mathcal{B}_0$ cannot be clearly interpreted as reversions *sensu stricto* along the terminal branch of *Chlorocebus sabaeus*. Indeed, we acknowledge that on a fixed fitness landscape, a deleterious mutation can be compensated by transitions to other fitter amino-acids, and not necessarily the ancestral one.

In the figures below, the first line is the name of the gene, the second line the position of the mutation in the OrthoMam protein-coding DNA alignment, and the arrow in the third line corresponds to the nucleotide position of the mutation. Finally, for reproducibility, we generated the fasta alignment for all $\mathcal{B}_0$ mutations that we detected, either in the terminal lineage or in segregating polymorphism. These fasta files are available at Zenodo (https://doi.org/10.5281/zenodo.7878953), across all populations, under the compressed archive *alignment_focused_around_non_adaptive_mutations.zip* and we provide a python script to obtain the figures shown below given these alignments.

Figure S2: Examples sites of $\mathcal{B}_0$ mutations in *Chlorocebus sabaeus* (reversions).

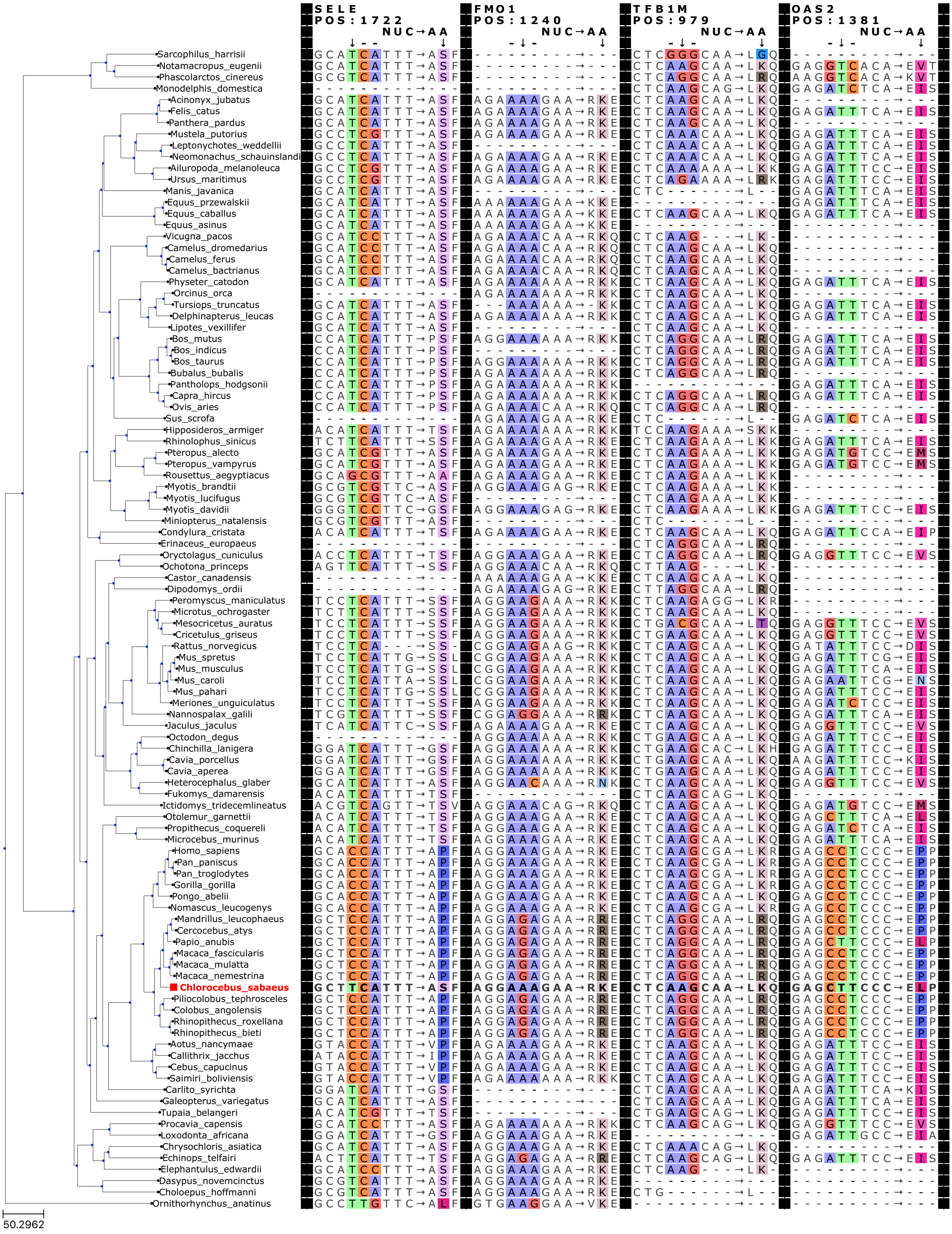

.

Figure S3: Examples sites of $\mathcal{B}_0$ mutations in *Chlorocebus sabaeus*.

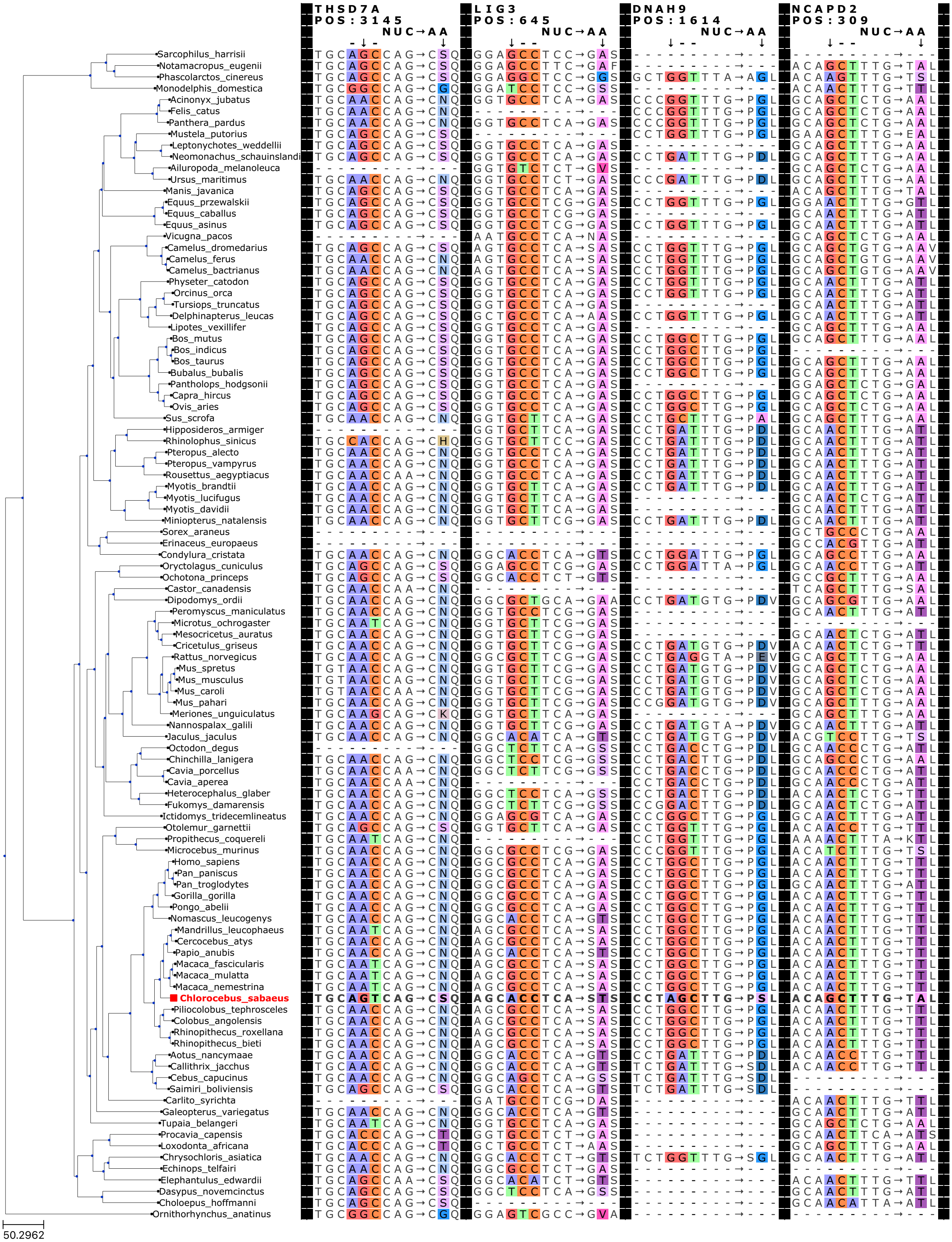

.

#### 2.2 Selection along the terminal branches

##### 2.2.1 Probability of mutations and substitutions to be $\mathcal{D}_0$, $\mathcal{N}_0$ or $\mathcal{B}_0$

Table S1: Probability of mutations and substitutions to be $\mathcal{D}_0$, $\mathcal{N}_0$ or $\mathcal{B}_0$.

| Population | Species | $\mathbb{P}[\mathcal{D}_0]$ | $\mathbb{P}[\mathcal{N}_0]$ | $\mathbb{P}[\mathcal{B}_0]$ | $\mathbb{P}_{div}[\mathcal{D}_0]$ | $\mathbb{P}_{div}[\mathcal{N}_0]$ | $\mathbb{P}_{div}[\mathcal{B}_0]$ |
|---|---|---|---|---|---|---|---|
| Equus c. | Equus caballus | 0.923 | 0.065 | 0.012 | 0.462 | 0.419 | 0.118 |
| Iran | Bos taurus | 0.924 | 0.065 | 0.011 | 0.515 | 0.362 | 0.123 |
| Uganda | Bos taurus | 0.924 | 0.065 | 0.011 | 0.514 | 0.361 | 0.125 |
| Australia | Capra hircus | 0.923 | 0.066 | 0.011 | 0.494 | 0.386 | 0.121 |
| France | Capra hircus | 0.923 | 0.066 | 0.011 | 0.494 | 0.386 | 0.120 |
| Iran (C. aegagrus) | Capra hircus | 0.923 | 0.066 | 0.011 | 0.493 | 0.386 | 0.120 |
| Iran | Capra hircus | 0.923 | 0.066 | 0.011 | 0.492 | 0.387 | 0.121 |
| Italy | Capra hircus | 0.923 | 0.066 | 0.011 | 0.494 | 0.386 | 0.120 |
| Morocco | Capra hircus | 0.923 | 0.066 | 0.011 | 0.491 | 0.387 | 0.122 |
| Iran | Ovis aries | 0.922 | 0.067 | 0.012 | 0.568 | 0.323 | 0.109 |
| Iran (O. orientalis) | Ovis aries | 0.922 | 0.067 | 0.011 | 0.573 | 0.320 | 0.108 |
| Iran (O. vignei) | Ovis aries | 0.922 | 0.067 | 0.012 | 0.567 | 0.325 | 0.109 |
| Various | Ovis aries | 0.922 | 0.067 | 0.011 | 0.572 | 0.321 | 0.107 |
| Morocco | Ovis aries | 0.922 | 0.067 | 0.012 | 0.570 | 0.321 | 0.108 |
| Barbados | Chlorocebus sabaeus | 0.926 | 0.065 | 0.009 | 0.485 | 0.393 | 0.122 |
| Central Afr. Rep. | Chlorocebus sabaeus | 0.926 | 0.065 | 0.009 | 0.485 | 0.391 | 0.124 |
| Ethiopia | Chlorocebus sabaeus | 0.926 | 0.065 | 0.009 | 0.484 | 0.393 | 0.124 |
| Gambia | Chlorocebus sabaeus | 0.926 | 0.065 | 0.009 | 0.483 | 0.394 | 0.123 |
| Kenya | Chlorocebus sabaeus | 0.926 | 0.065 | 0.009 | 0.485 | 0.392 | 0.123 |
| Nevis | Chlorocebus sabaeus | 0.926 | 0.065 | 0.009 | 0.484 | 0.393 | 0.123 |
| South Africa | Chlorocebus sabaeus | 0.926 | 0.065 | 0.009 | 0.480 | 0.394 | 0.125 |
| Saint Kitts | Chlorocebus sabaeus | 0.926 | 0.065 | 0.009 | 0.483 | 0.394 | 0.123 |
| Zambia | Chlorocebus sabaeus | 0.926 | 0.065 | 0.009 | 0.485 | 0.393 | 0.123 |
| African | Homo sapiens | 0.925 | 0.065 | 0.010 | 0.561 | 0.341 | 0.099 |
| Admixed American | Homo sapiens | 0.925 | 0.065 | 0.010 | 0.561 | 0.340 | 0.099 |
| East Asian | Homo sapiens | 0.925 | 0.065 | 0.010 | 0.560 | 0.341 | 0.098 |
| European | Homo sapiens | 0.925 | 0.065 | 0.010 | 0.562 | 0.340 | 0.098 |
| South Asian | Homo sapiens | 0.925 | 0.065 | 0.010 | 0.561 | 0.341 | 0.099 |

- $\mathbb{P}[\mathcal{D}_0]$ (eq. 5) is the probability for a new mutation to be a deleterious. These mutations have a selection coefficient predicted at the phylogenetic-scale lower than -1, thus toward a less fit amino-acid.
- $\mathbb{P}[\mathcal{N}_0]$ (eq. 5) is the probability for a new mutation to be a nearly-neutral. These mutations have a selection coefficient predicted at the phylogenetic-scale between -1 and 1.
- $\mathbb{P}[\mathcal{B}_0]$ (eq. 5) is the probability for a new mutation to be a non-adaptive beneficial mutation. These mutations have a selection coefficient predicted at the phylogenetic-scale larger than 1, thus toward a more fit amino-acid.
- $\mathbb{P}_{div}[\mathcal{D}_0]$ is the proportion of substitutions in the terminal branch that are $\mathcal{D}_0$.
- $\mathbb{P}_{div}[\mathcal{N}_0]$ is the proportion of substitutions in the terminal branch that are $\mathcal{N}_0$.
- $\mathbb{P}_{div}[\mathcal{B}_0]$ is the proportion of substitutions in the terminal branch that are $\mathcal{B}_0$.

##### 2.2.2 $d_N/d_S$ for $\mathcal{D}_0$, $\mathcal{N}_0$ or $\mathcal{B}_0$

Table S2: $d_N/d_S$ for $\mathcal{D}_0$, $\mathcal{N}_0$ or $\mathcal{B}_0$.

| Species | $d_N/d_S$ | $d_N(\mathcal{D}_0)/d_S$ | $d_N(\mathcal{N}_0)/d_S$ | $d_N(\mathcal{B}_0)/d_S$ |
|---|---|---|---|---|
| Equus caballus | 0.129 | 0.065 | 0.832 | 1.267 |
| Bos taurus | [0.114, 0.116] | [0.063, 0.064] | [0.629, 0.638] | [1.280, 1.328] |
| Capra hircus | [0.108, 0.109] | [0.058, 0.058] | [0.631, 0.636] | [1.169, 1.183] |
| Ovis aries | [0.127, 0.129] | [0.078, 0.080] | [0.619, 0.621] | [1.201, 1.217] |
| Chlorocebus sabaeus | [0.118, 0.119] | [0.062, 0.062] | [0.713, 0.720] | [1.521, 1.577] |
| Homo sapiens | [0.170, 0.170] | [0.103, 0.103] | [0.884, 0.888] | [1.733, 1.745] |

- $d_N/d_S$ (eq. 6) is the ratio of non-synonymous over synonymous substitutions estimated for all the non-synonymous substitutions in the terminal branch.
- $d_N(\mathcal{D}_0)/d_S$ (eq. 6) is the ratio of non-synonymous over synonymous substitutions, when restricted to non-synonymous substitutions in the terminal branch that are $\mathcal{D}_0$.
- $d_N(\mathcal{N}_0)/d_S$ (eq. 6) is the ratio of non-synonymous over synonymous substitutions, when restricted to non-synonymous substitutions in the terminal branch that are $\mathcal{N}_0$.
- $d_N(\mathcal{B}_0)/d_S$ (eq. 6) is the ratio of non-synonymous over synonymous substitutions, when restricted to non-synonymous substitutions in the terminal branch that are $\mathcal{B}_0$.

SNP are considered fixed in the populations if all sampled individuals are homozygous for the derived allele, in which case we consider this SNP not to be segregating but as fixed as a substitution. This effect results in $d_N/d_S$ that can change across populations, denoted as a range per species. $d_N(\mathcal{B}_0)/d_S$ values observed are above 1 and are consistent with slightly advantageous mutations.

Theoretically, $\omega = d_N/d_S$ can be related to the underlying scaled selection coefficient ($S$) with the relation $\omega = S/(1 - \exp(-S))$ as in [6, eq. 3]. In our experiments our observed $d_N(\mathcal{B}_0)/d_S$ is on the range 1.169 (*C. hircus*) to 1.745 (*H. sapiens*), translating into average $S$ of $\approx 0.32$ to $\approx 1.24$, so indeed only slightly advantageous.

Of note, observing $d_N(\mathcal{B}_0)/d_S > 1$ is an important check that $\mathcal{B}_0$ mutations are indeed positively selected with an increased substitution rate. Since $d_N(\mathcal{N}_0)/d_S$ is close to 1 for the predicted nearly-neutral mutations, this means that the selection coefficients predicted at the mutation-selection balance are good proxies of selection, then observing $d_N(\mathcal{B}_0)/d_S > 1$ is evidence of positive selection of predicted non-adaptive beneficial mutations.

##### 2.2.3 $d_N/d_S$ over-estimation due to non-adaptive beneficial mutations

Table S3: $d_N/d_S$ over-estimation due to non-adaptive beneficial mutations.

| Species | $d_N/d_S$ | $d_N(S_0 < 1)/d_S$ | $\delta(d_N/d_S)$ |
|---|---|---|---|
| Equus caballus | 0.129 | 0.115 | 10.7 |
| Bos taurus | [0.114, 0.116] | [0.101, 0.102] | [11.3, 11.5] |
| Capra hircus | [0.108, 0.109] | [0.096, 0.097] | [11.0, 11.2] |
| Chlorocebus sabaeus | [0.118, 0.119] | [0.104, 0.105] | [11.4, 11.7] |
| Homo sapiens | [0.170, 0.170] | [0.154, 0.155] | [8.926, 8.996] |
| Ovis aries | [0.127, 0.129] | [0.115, 0.117] | [9.663, 9.894] |

- $d_N/d_S$ (eq. 6) is the ratio of non-synonymous over synonymous substitutions estimated for all the non-synonymous substitutions in the terminal branch.
- $d_N(S_0 < 1)/d_S$ (eq. 6) is the ratio of non-synonymous over synonymous substitutions, when restricted to non-synonymous substitutions in the terminal branch that are not $\mathcal{B}_0$. This is the estimated divergence when we removed non-adaptive beneficial mutations.
- $\delta(d_N/d_S)$ (eq. 7) is the fraction of the divergence ($d_N/d_S$) that is over-estimated, $d_N/d_S$ is compared to the estimated divergence when we removed non-adaptive beneficial mutations $d_N(S_0 < 1)/d_S$.

We estimated that between 9 and 11% of $d_N/d_S$ is over-estimated, corresponding to non-adaptive beneficial mutations inflating the $d_N/d_S$ statistic.

### 3 Gene ontology enrichment

#### 3.1 Gene ontology enrichment in $\mathcal{B}_0$ SNPs

To assess whether $\mathcal{B}_0$ SNPs are enriched in specific gene functions, we used gene ontology (GO) terms at the gene level. Thus, for each $\mathcal{B}_0$ SNP detected in a focal population, we annotated this SNP with GO terms inherited from the gene in which the SNP is located. We restricted this analysis solely on genes annotated with at least one GO term.

In order to assess the weight of each GO term in the set of $\mathcal{B}_0$ SNPs, we computed the proportion of $\mathcal{B}_0$ SNPs annotated with a given GO term. Thus, for each GO term, $p$ is the proportion of $\mathcal{B}_0$ SNPs that are annotated with this focal GO term among all annotated $\mathcal{B}_0$ SNPs.

In order to test if the distribution of $\mathcal{B}_0$ SNPs is different between genes sharing a common GO term and the rest of the genes, we used a Mann-Whitney U test. To do so, first, for each gene, we computed the proportion of sites that have a $\mathcal{B}_0$ SNP, called $p[\mathcal{B}_0]$ (can be 0 if no $\mathcal{B}_0$ SNP is detected for this gene). Second, for each GO term, $r$ is the ratio of the mean of $p[\mathcal{B}_0]$ in the set of genes sharing the focal ontology over the mean of $p[\mathcal{B}_0]$ in the rest of the genes. Third, Mann-Whitney U test (two-sided) is used to test if the distribution of $p[\mathcal{B}_0]$ is different between the two sets of genes (U statistic and p-value $p_\text{v}$). Finally, $p_\text{v}^\text{adj}$ are corrected for multiple comparison (Holm–Bonferroni correction across the ontology terms). * for $p_\text{v}^\text{adj}$ corrected for multiple comparison (Holm–Bonferroni correction) lower than the risk $\alpha = 0.05$.

The table below shows the GO terms (first 100 rows with the lowest $p_\text{v}$) that are enriched in genes with $\mathcal{B}_0$ SNP in human (european population).

Table S4: Gene ontology enrichment in $\mathcal{B}_0$ SNPs

| GO id | GO name | $p$ | $r$ | Mann-Whitney U | $p_\text{v}$ | $p_\text{v}^\text{adj}$ |
|---|---|---|---|---|---|---|
| GO:0003777 | microtubule motor activity | 0.027 | 1.802 | $1.1 \times 10^5$ | $1.8 \times 10^{-6}$ | **0.0008*** |
| GO:0005578 | proteinaceous extracellular matrix | 0.069 | 2.071 | $4.7 \times 10^5$ | $1.9 \times 10^{-6}$ | **0.00084*** |
| GO:0003774 | motor activity | 0.023 | 1.127 | $1.1 \times 10^5$ | $6.7 \times 10^{-5}$ | **0.029*** |
| GO:0030198 | extracellular matrix organization | 0.039 | 1.956 | $2.7 \times 10^5$ | 0.00055 | 0.239 |
| GO:0007018 | microtubule-based movement | 0.023 | 0.924 | $1.3 \times 10^5$ | 0.00073 | 0.315 |
| GO:0005576 | extracellular region | 0.178 | 2.195 | $1.9 \times 10^6$ | 0.00083 | 0.359 |
| GO:0006898 | receptor-mediated endocytosis | 0.027 | 1.803 | $1.7 \times 10^5$ | 0.001 | 0.512 |
| GO:0006355 | regulation of transcription | 0.069 | 0.275 | $1.9 \times 10^6$ | 0.002 | 0.815 |
| GO:0005604 | basement membrane | 0.019 | 1.638 | $1.1 \times 10^5$ | 0.002 | 0.852 |
| GO:0007229 | integrin-mediated signaling pathway | 0.019 | 1.930 | $1.1 \times 10^5$ | 0.003 | 1.000 |
| GO:0016887 | ATPase activity | 0.031 | 1.149 | $1.8 \times 10^5$ | 0.003 | 1.000 |
| GO:0007155 | cell adhesion | 0.062 | 1.438 | $5.8 \times 10^5$ | 0.005 | 1.000 |
| GO:0005634 | nucleus | 0.266 | 0.557 | $3.9 \times 10^6$ | 0.005 | 1.000 |
| GO:0006351 | transcription | 0.073 | 0.339 | $1.9 \times 10^6$ | 0.008 | 1.000 |
| GO:0005654 | nucleoplasm | 0.124 | 0.443 | $2.6 \times 10^6$ | 0.009 | 1.000 |
| GO:0006366 | transcription from RNA polymerase II promoter | 0.008 | 0.063 | $6.2 \times 10^5$ | 0.011 | 1.000 |
| GO:0005829 | cytosol | 0.212 | 0.541 | $3.5 \times 10^6$ | 0.012 | 1.000 |
| GO:0016853 | isomerase activity | 0.015 | 4.066 | $1 \times 10^5$ | 0.015 | 1.000 |
| GO:0071356 | cellular response to tumor necrosis factor | 0.015 | 7.069 | $1 \times 10^5$ | 0.016 | 1.000 |
| GO:0005518 | collagen binding | 0.015 | 2.900 | $1 \times 10^5$ | 0.017 | 1.000 |
| GO:0043087 | regulation of GTPase activity | 0.015 | 2.025 | $1.1 \times 10^5$ | 0.027 | 1.000 |
| GO:0000139 | Golgi membrane | 0.012 | 0.214 | $6.3 \times 10^5$ | 0.028 | 1.000 |
| GO:0043235 | receptor complex | 0.023 | 0.935 | $1.9 \times 10^5$ | 0.028 | 1.000 |
| GO:0000981 | RNA polymerase II transcription factor activity | 0.027 | 0.452 | $9.1 \times 10^5$ | 0.029 | 1.000 |
| GO:0005739 | mitochondrion | 0.058 | 0.471 | $1.4 \times 10^6$ | 0.029 | 1.000 |
| GO:0005581 | collagen trimer | 0.015 | 2.124 | $1.1 \times 10^5$ | 0.036 | 1.000 |
| GO:0051015 | actin filament binding | 0.019 | 0.912 | $1.5 \times 10^5$ | 0.036 | 1.000 |
| GO:0001843 | neural tube closure | 0.015 | 1.164 | $1.1 \times 10^5$ | 0.036 | 1.000 |

Continued on next page

| GO id | GO name | $p$ | $r$ | Mann-Whitney U | $p_v$ | $p_v^{adj}$ |
|---|---|---|---|---|---|---|
| GO:0005737 | cytoplasm | 0.317 | 0.692 | $4.1 \times 10^6$ | 0.037 | 1.000 |
| GO:0043565 | sequence-specific DNA binding | 0.015 | 0.308 | $6 \times 10^5$ | 0.039 | 1.000 |
| GO:0004222 | metalloendopeptidase activity | 0.019 | 1.188 | $1.6 \times 10^5$ | 0.043 | 1.000 |
| GO:0007166 | cell surface receptor signaling pathway | 0.031 | 2.871 | $3 \times 10^5$ | 0.046 | 1.000 |
| GO:0007156 | homophilic cell adhesion via plasma membrane... | 0.015 | 1.013 | $1.2 \times 10^5$ | 0.050 | 1.000 |
| GO:0007286 | spermatid development | 0.015 | 2.497 | $1.2 \times 10^5$ | 0.051 | 1.000 |
| GO:0008202 | steroid metabolic process | 0.015 | 2.755 | $1.2 \times 10^5$ | 0.051 | 1.000 |
| GO:0061024 | membrane organization | 0.019 | 1.516 | $1.6 \times 10^5$ | 0.052 | 1.000 |
| GO:1901796 | regulation of signal transduction by p53 class... | 0.019 | 1.333 | $1.6 \times 10^5$ | 0.058 | 1.000 |
| GO:0006397 | mRNA processing | 0.004 | 0.210 | $3.5 \times 10^5$ | 0.061 | 1.000 |
| GO:0006915 | apoptotic process | 0.015 | 0.463 | $6.4 \times 10^5$ | 0.065 | 1.000 |
| GO:0043547 | positive regulation of GTPase activity | 0.035 | 2.651 | $3.7 \times 10^5$ | 0.068 | 1.000 |
| GO:0007010 | cytoskeleton organization | 0.019 | 1.771 | $1.7 \times 10^5$ | 0.068 | 1.000 |
| GO:0030308 | negative regulation of cell growth | 0.015 | 3.055 | $1.3 \times 10^5$ | 0.070 | 1.000 |
| GO:0007160 | cell-matrix adhesion | 0.015 | 1.046 | $1.2 \times 10^5$ | 0.073 | 1.000 |
| GO:0006897 | endocytosis | 0.023 | 1.408 | $2.2 \times 10^5$ | 0.073 | 1.000 |
| GO:0003677 | DNA binding | 0.077 | 0.448 | $1.7 \times 10^6$ | 0.075 | 1.000 |
| GO:0043202 | lysosomal lumen | 0.015 | 1.522 | $1.3 \times 10^5$ | 0.075 | 1.000 |
| GO:0003700 | DNA binding transcription factor activity | 0.027 | 0.343 | $8.7 \times 10^5$ | 0.076 | 1.000 |
| GO:0008134 | transcription factor binding | 0.004 | 0.048 | $3.2 \times 10^5$ | 0.077 | 1.000 |
| GO:0005488 | binding | 0.046 | 0.641 | $4.1 \times 10^5$ | 0.078 | 1.000 |
| GO:0045944 | positive regulation of transcription from RNA... | 0.039 | 0.324 | $1.1 \times 10^6$ | 0.085 | 1.000 |
| GO:0005794 | Golgi apparatus | 0.046 | 0.430 | $1.2 \times 10^6$ | 0.087 | 1.000 |
| GO:0016032 | viral process | 0.008 | 0.321 | $4.1 \times 10^5$ | 0.089 | 1.000 |
| GO:0055085 | transmembrane transport | 0.054 | 1.533 | $5.9 \times 10^5$ | 0.092 | 1.000 |
| GO:0006260 | DNA replication | 0.019 | 1.051 | $1.8 \times 10^5$ | 0.096 | 1.000 |
| GO:0030425 | dendrite | 0.008 | 0.121 | $3.9 \times 10^5$ | 0.099 | 1.000 |
| GO:0005524 | ATP binding | 0.127 | 0.666 | $1.5 \times 10^6$ | 0.103 | 1.000 |
| GO:0005615 | extracellular space | 0.100 | 1.699 | $1.3 \times 10^6$ | 0.107 | 1.000 |
| GO:0007283 | spermatogenesis | 0.039 | 1.824 | $3.9 \times 10^5$ | 0.108 | 1.000 |
| GO:0005179 | hormone activity | 0.012 | 7.683 | $9.9 \times 10^4$ | 0.118 | 1.000 |
| GO:0098793 | presynapse | 0.012 | 2.065 | $9.6 \times 10^4$ | 0.119 | 1.000 |
| GO:0005254 | chloride channel activity | 0.012 | 1.694 | $9.6 \times 10^4$ | 0.124 | 1.000 |
| GO:0036064 | ciliary basal body | 0.015 | 1.378 | $1.4 \times 10^5$ | 0.125 | 1.000 |
| GO:0070062 | extracellular exosome | 0.104 | 0.570 | $2.1 \times 10^6$ | 0.125 | 1.000 |
| GO:0090630 | activation of GTPase activity | 0.012 | 2.519 | $9.9 \times 10^4$ | 0.127 | 1.000 |
| GO:0044212 | transcription regulatory region DNA binding | 0.004 | 0.322 | $2.7 \times 10^5$ | 0.128 | 1.000 |
| GO:0055114 | oxidation-reduction process | 0.019 | 0.482 | $6.4 \times 10^5$ | 0.129 | 1.000 |
| GO:0019886 | antigen processing and presentation of exogenous... | 0.012 | 1.393 | $9.6 \times 10^4$ | 0.129 | 1.000 |
| GO:0005743 | mitochondrial inner membrane | 0.008 | 0.463 | $3.7 \times 10^5$ | 0.133 | 1.000 |
| GO:0022617 | extracellular matrix disassembly | 0.012 | 1.009 | $9.6 \times 10^4$ | 0.135 | 1.000 |
| GO:0042493 | response to drug | 0.004 | 0.043 | $2.6 \times 10^5$ | 0.137 | 1.000 |
| GO:0008380 | RNA splicing | 0.008 | 0.098 | $2.6 \times 10^5$ | 0.139 | 1.000 |
| GO:0043687 | post-translational protein modification | 0.008 | 0.319 | $3.6 \times 10^5$ | 0.140 | 1.000 |
| GO:0010951 | negative regulation of endopeptidase activity | 0.012 | 2.037 | $9.9 \times 10^4$ | 0.140 | 1.000 |
| GO:0000978 | RNA polymerase II proximal promoter... | 0.012 | 0.250 | $4.5 \times 10^5$ | 0.140 | 1.000 |
| GO:0003713 | transcription coactivator activity | 0.004 | 0.339 | $2.6 \times 10^5$ | 0.142 | 1.000 |
| GO:0001227 | transcriptional repressor activity | 0.012 | 1.384 | $9.9 \times 10^4$ | 0.144 | 1.000 |
| GO:0005623 | cell | 0.012 | 1.927 | $1 \times 10^5$ | 0.150 | 1.000 |
| GO:0030574 | collagen catabolic process | 0.012 | 1.036 | $9.9 \times 10^4$ | 0.151 | 1.000 |
| GO:0006457 | protein folding | 0.015 | 2.465 | $1.5 \times 10^5$ | 0.158 | 1.000 |
| GO:0016477 | cell migration | 0.004 | 0.121 | $2.4 \times 10^5$ | 0.159 | 1.000 |
| GO:0070374 | positive regulation of ERK1 and ERK2 cascade | 0.019 | 4.238 | $2 \times 10^5$ | 0.160 | 1.000 |
| GO:0007605 | sensory perception of sound | 0.015 | 0.824 | $1.5 \times 10^5$ | 0.162 | 1.000 |
| GO:0031225 | anchored component of membrane | 0.012 | 0.892 | $1 \times 10^5$ | 0.171 | 1.000 |
| GO:0016757 | transferase activity | 0.004 | 0.435 | $2.4 \times 10^5$ | 0.177 | 1.000 |
| GO:0002376 | immune system process | 0.012 | 0.411 | $4.3 \times 10^5$ | 0.177 | 1.000 |
| GO:0005759 | mitochondrial matrix | 0.008 | 0.361 | $3.3 \times 10^5$ | 0.185 | 1.000 |
| GO:0010389 | regulation of G2/M transition of mitotic cell... | 0.012 | 0.924 | $1 \times 10^5$ | 0.188 | 1.000 |
| GO:0005667 | transcription factor complex | 0.004 | 0.177 | $2.3 \times 10^5$ | 0.189 | 1.000 |
| GO:0030054 | cell junction | 0.031 | 0.251 | $8.1 \times 10^5$ | 0.190 | 1.000 |
| GO:0001650 | fibrillar center | 0.019 | 1.540 | $1.1 \times 10^5$ | 0.191 | 1.000 |
| GO:0014069 | postsynaptic density | 0.004 | 0.172 | $2.2 \times 10^5$ | 0.200 | 1.000 |
| GO:0046934 | phosphatidylinositol-4 | 0.012 | 1.135 | $1.1 \times 10^5$ | 0.201 | 1.000 |
| GO:0007267 | cell-cell signaling | 0.004 | 0.354 | $2.2 \times 10^5$ | 0.201 | 1.000 |
| GO:0001649 | osteoblast differentiation | 0.012 | 1.917 | $1.1 \times 10^5$ | 0.201 | 1.000 |
| GO:0045893 | positive regulation of transcription | 0.023 | 0.596 | $6.6 \times 10^5$ | 0.205 | 1.000 |
| GO:0007399 | nervous system development | 0.015 | 0.450 | $5 \times 10^5$ | 0.205 | 1.000 |
| GO:0001501 | skeletal system development | 0.015 | 1.212 | $1.6 \times 10^5$ | 0.207 | 1.000 |
| GO:0019903 | protein phosphatase binding | 0.012 | 0.875 | $1.1 \times 10^5$ | 0.208 | 1.000 |
| GO:1902476 | chloride transmembrane transport | 0.012 | 1.458 | $1.1 \times 10^5$ | 0.210 | 1.000 |
| GO:0007568 | aging | 0.015 | 2.053 | $1.6 \times 10^5$ | 0.210 | 1.000 |

#### 3.2 Enrichment in $\mathcal{B}_0$ SNPs across all gene ontology terms

Across all 347 gene ontology terms, the distribution of $p_{\text{v}}$ is shown in the histogram below.

Figure S4: Enrichment in $\mathcal{B}_0$ SNPs across all gene ontology terms.

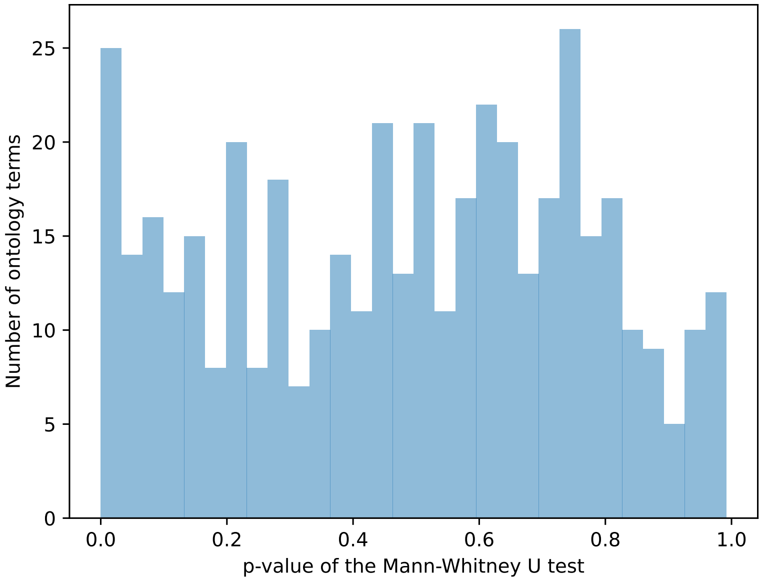

### 4 Clinically related terms for mutations

#### 4.1 Terms associated with deleterious mutations $\mathcal{D}_0$

Table S5: Terms associated with deleterious mutations $\mathcal{D}_0$.

| SNP clinical ontology | $n_{\text{Observed}}$ | $n_{\text{Expected}}$ | Odds ratio | $p_{\text{v}}$ | $p_{\text{v-adjusted}}$ |
|---|---|---|---|---|---|
| Benign | 2969 | 4043.0 | 0.734 | 1.000 | 1.000 |
| Likely benign | 2994 | 3399.8 | 0.881 | 0.999 | 1.000 |
| Risk factor | 102 | 118.2 | 0.863 | 0.798 | 1.000 |
| Likely pathogenic | 221 | 68.5 | 3.226 | $1.7 \times 10^{-8}$ | $\mathbf{6.7 \times 10^{-8}}$* |
| Pathogenic | 560 | 193.6 | 2.893 | $4.2 \times 10^{-17}$ | $\mathbf{2.1 \times 10^{-16}}$* |

In humans (european population), non-synonymous SNPs in the test group ($\mathcal{D}_0$) are contrasted to SNPs in the control group ($\mathcal{N}_0$). For each clinical term, a 2x2 contingency tables is built by counting the number of SNPs based on their selection coefficient and their clinical terms (whether they have this specific term or not). Fisher's exact tests are then performed for these 2x2 contingency tables. * for $p_{\text{v}}^{\text{adj}}$ corrected for multiple comparison (Holm–Bonferroni correction) lower than the risk $\alpha = 0.05$. SNPs predicted with $\mathcal{D}_0$ are statistically associated to clinical terms such as *Likely Pathogenic* and *Pathogenic*.

#### 4.2 Terms associated with non-adaptive beneficial mutations $\mathcal{B}_0$

Table S6: Terms associated with non-adaptive beneficial mutations $\mathcal{B}_0$.

| SNP clinical ontology | $n_{\text{Observed}}$ | $n_{\text{Expected}}$ | Odds ratio | $p_{\text{v}}$ | $p_{\text{v-adjusted}}$ |
|---|---|---|---|---|---|
| Benign | 319 | 261.7 | 1.219 | 0.002 | **0.009*** |
| Likely benign | 263 | 222.7 | 1.181 | 0.012 | **0.049*** |
| Risk factor | 5 | 7.847 | 0.637 | 0.879 | 0.879 |
| Likely pathogenic | 7 | 4.552 | 1.538 | 0.227 | 0.682 |
| Pathogenic | 16 | 12.9 | 1.241 | 0.268 | 0.682 |

In humans, non-synonymous SNPs in the test group ($\mathcal{B}_0$) are contrasted to SNPs in the control group ($\mathcal{N}_0$). For each clinical term, a 2x2 contingency tables is built by counting the number of SNPs based on their selection coefficient and their clinical terms (whether they have this specific term or not). Fisher's exact tests are then performed for these 2x2 contingency tables. * for $p_{\text{v}}^{\text{adj}}$ corrected for multiple comparison (Holm–Bonferroni correction) lower than the risk $\alpha = 0.05$. Beneficial non-adaptive mutations are associated with clinical terms such as *Benign* and *Likely Benign*.

### 5 Correlation with diversity

#### 5.1 Phylogenetic Generalized Linear Model

Because a correlation must account for phylogenetic relationship and non-independence of samples, we fitted a Phylogenetic Generalized Linear Model (PGLM) in R with the package caper[7], with multi-furcation of the different populations inside each species. The mammalian tree imported from TimeTree[8] and pruned to the species used in this study is, in newick format:

```
((Equus:76,((IRBT:3.55912,UGBT:3.55912)1993:18.0665,((IROA:5.33074,IROO:5.33074,ISGC
    :5.33074,IROV:5.33074,MOOA:5.33074)2219:2.32334,(IRCA:3.96087,IRCH:3.96087,AUCH
    :3.96087,FRCH:3.96087,ITCH:3.96087,MOCH:3.96087)2201:3.69321)2218:13.9716)
    2262:54.3744)2405:18,((AFR:5.59333,AMR:5.59333,EAS:5.59333,EUR:5.59333,SAS
    :5.59333)8454:22.42,(Barbados:2.69204,CAR:2.69204,Ethiopia:2.69204,Gambia:2.69204,
    Kenya:2.69204,Nevis:2.69204,SK:2.69204,SA:2.69204,Zambia:2.69204)8449:26.128)
    8800:65.18);
```

And shown in the figure below:

Figure S5: Tree used for PGLM (Phylogenetic Generalized Linear Model).

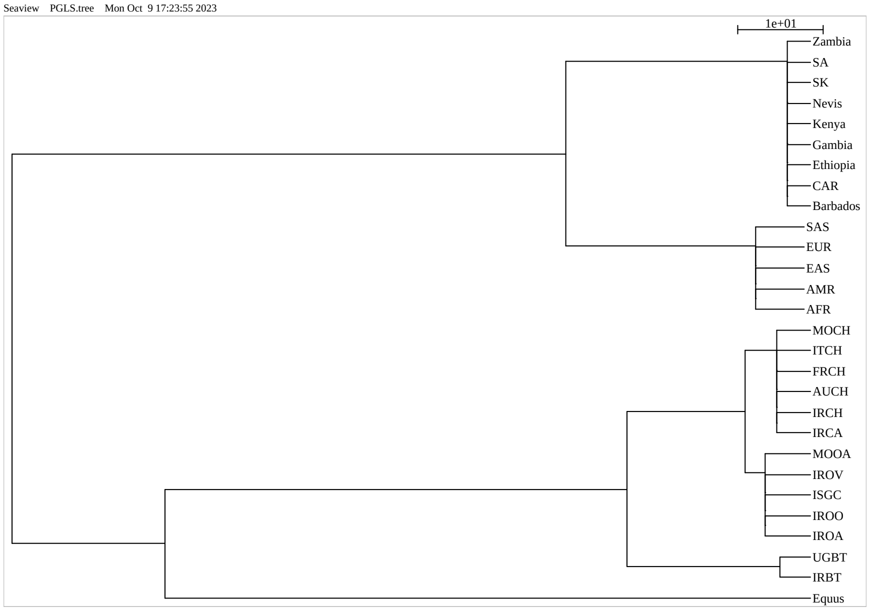

Multifurcations of the different populations are placed at the same divergence time as the species. Then, for each population, the proportion of deleterious ($\mathbb{P}[\mathcal{D}]$), nearly-neutral ($\mathbb{P}[\mathcal{N}]$) and of beneficial ($\mathbb{P}[\mathcal{B}]$) mutations estimated at the population-genetic scale is shown as function of $N_{\mathrm{e}}$. $r^2$ and p-value are obtained from the PGLM model.

##### 5.1.1 Proportion of deleterious mutations ($\mathcal{D}$)

Figure S6: Proportion of deleterious mutations ($\mathcal{D}$).

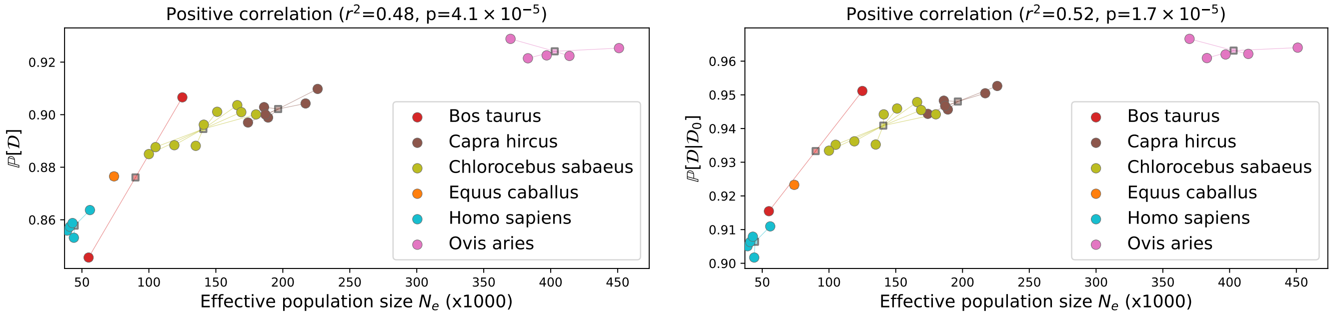

- $N_\mathrm{e}$ is the estimated effective population size.
- $\mathbb{P}[\mathcal{D}]$ (eq. 15) is the probability for a mutation to be deleterious. These mutations have a selection coefficient at the population-scale lower than -1.
- $\mathbb{P}[\mathcal{D} \mid \mathcal{D}_0]$ (eq. 12) is the probability for a mutation to be deleterious at the population scale, given it is predicted to be a deleterious at the phylogenetic scale.

We can see that higher effective population size ($N_\mathrm{e}$) is typically accompanied by a higher proportion of effectively deleterious mutations at the population scale ($\mathbb{P}[\mathcal{D}]$). This trend is also confirmed when we restricted the analysis to class of mutations that are supposedly deleterious at the phylogenetic scale ($\mathcal{D}_0$). This result is qualitatively in accordance with the nearly-neutral theory of evolution which argues that very slightly deleterious mutations are more efficiently purified in large populations.

##### 5.1.2 Proportion of nearly-neutral mutations ($\mathcal{N}$)

Figure S7: Proportion of nearly-neutral mutations ($\mathcal{N}$).

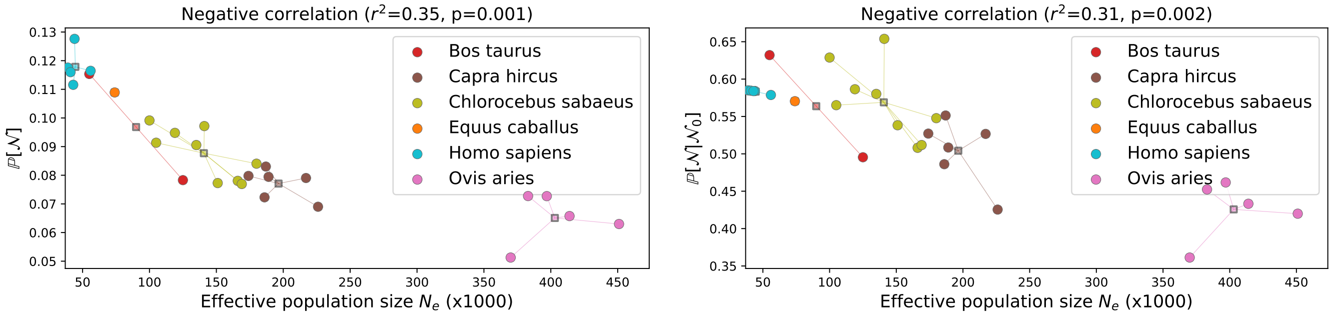

- $N_\mathrm{e}$ is the estimated effective population size.
- $\mathbb{P}[\mathcal{N}]$ (eq. 15) is the probability for a mutation to be nearly-neutral. These mutations have a selection coefficient at the population-scale between -1 and 1.
- $\mathbb{P}[\mathcal{N} \mid \mathcal{N}_0]$ (eq. 12) is the probability for a mutation to be nearly-neutral at the population scale, given it is predicted to be a nearly-neutral at the phylogenetic scale.

We confirmed that higher effective population size ($N_\mathrm{e}$) is typically accompanied by a smaller proportion of neutral mutations at the population scale ($\mathbb{P}[\mathcal{N}]$ in the range 0.06-0.18). This result is more pronounced ($\mathbb{P}[\mathcal{N} \mid \mathcal{N}_0]$ in the range 0.36-0.73) when we restricted the analysis to class of mutations that are supposedly nearly-neutral at the phylogenetic scale ($\mathcal{N}_0$). This result suggests that populations with higher diversity (e.g. *Bos* or *Ovis*) are more likely to discriminate whether mutations are beneficial or deleterious. Alternatively stated, mutations in populations with low diversity (e.g. *Homo*) are effectively nearly-neutral and behave as a neutral mutation would. This result is qualitatively in accordance with the nearly-neutral theory of evolution which argues that mutations are less efficiently selected for in small populations.

##### 5.1.3 Proportion of beneficial mutations ($\mathcal{B}$)

Figure S8: Proportion of beneficial mutations ($\mathcal{B}$).

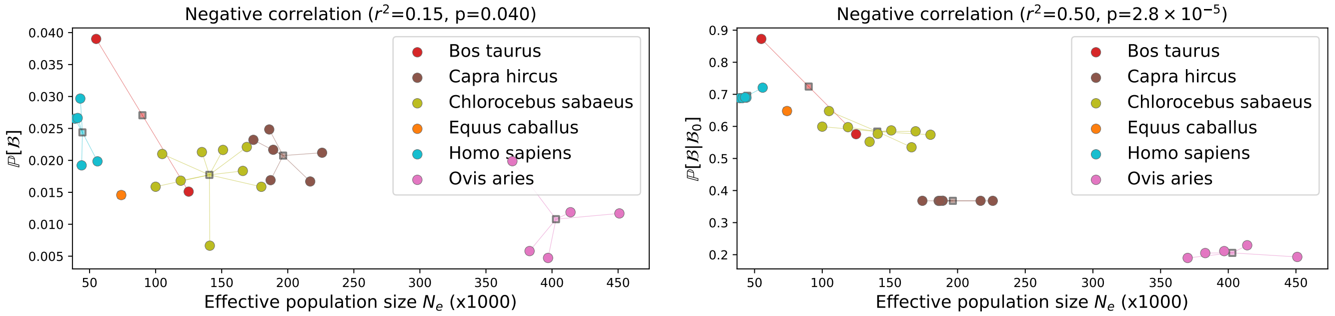

- $N_\text{e}$ is the estimated effective population size.
- $\mathbb{P}[\mathcal{B}]$ (eq. 15) is the probability for a mutation to be beneficial. These mutations have a selection coefficient at the population-scale larger than 1.
- $\mathbb{P}[\mathcal{B} \mid \mathcal{B}_0]$ (eq. 12) is the probability for a mutation to be beneficial at the population scale, given it is predicted to be a non-adaptive beneficial mutation at the phylogenetic scale (the *precision*).

Higher effective population size ($N_\text{e}$) is accompanied by a smaller proportion of beneficial mutations at the population scale ($\mathbb{P}[\mathcal{B}]$). This trend is also confirmed when we restricted the analysis to class of mutations that are supposedly non-adaptive beneficial mutations at the phylogenetic scale ($\mathcal{B}_0$).

### 6 Probability for beneficial mutations to be non-adaptive ($\mathbb{P}[\mathcal{B}_0 \mid \mathcal{B}]$)

#### 6.1 Across the different populations

Table S7: Probability for beneficial mutations to be non-adaptive ($\mathbb{P}[\mathcal{B}_0 \mid \mathcal{B}]$).

| Population | Species | $N_e$ | $\mathbb{P}[\mathcal{B}_0]$ | $\mathbb{P}[\mathcal{B}]$ | $\frac{\mathbb{P}[\mathcal{B}_0]}{\mathbb{P}[\mathcal{B}]}$ | $\mathbb{P}[\mathcal{B} \mid \mathcal{B}_0]$ | $\mathbb{P}[\mathcal{B}_0 \mid \mathcal{B}]$ |
|---|---|---|---|---|---|---|---|
| Equus c. | Equus caballus | $7.5 \times 10^4$ | 0.012 | 0.015 | 0.827 | 0.648 | 0.536 |
| Iran | Bos taurus | $5.6 \times 10^4$ | 0.011 | 0.039 | 0.279 | 0.873 | 0.243 |
| Uganda | Bos taurus | $1.3 \times 10^5$ | 0.011 | 0.015 | 0.720 | 0.576 | 0.415 |
| Australia | Capra hircus | $1.7 \times 10^5$ | 0.011 | 0.023 | 0.480 | 0.368 | 0.177 |
| France | Capra hircus | $1.9 \times 10^5$ | 0.011 | 0.022 | 0.515 | 0.368 | 0.190 |
| Iran (C. aegagrus) | Capra hircus | $1.9 \times 10^5$ | 0.011 | 0.025 | 0.448 | 0.368 | 0.165 |
| Iran | Capra hircus | $2.3 \times 10^5$ | 0.011 | 0.021 | 0.525 | 0.368 | 0.193 |
| Italy | Capra hircus | $1.9 \times 10^5$ | 0.011 | 0.017 | 0.660 | 0.368 | 0.243 |
| Morocco | Capra hircus | $2.2 \times 10^5$ | 0.011 | 0.017 | 0.667 | 0.368 | 0.245 |
| Iran | Ovis aries | $3.8 \times 10^5$ | 0.012 | 0.006 | 1.984 | 0.205 | 0.407 |
| Iran (O. orientalis) | Ovis aries | $4.5 \times 10^5$ | 0.011 | 0.012 | 0.983 | 0.193 | 0.190 |
| Iran (O. vignei) | Ovis aries | $3.7 \times 10^5$ | 0.012 | 0.020 | 0.579 | 0.190 | 0.110 |
| Various | Ovis aries | $4.1 \times 10^5$ | 0.011 | 0.012 | 0.967 | 0.229 | 0.222 |
| Morocco | Ovis aries | $4 \times 10^5$ | 0.012 | 0.005 | 2.435 | 0.211 | 0.514 |
| Barbados | Chlorocebus sabaeus | $1.1 \times 10^5$ | 0.009 | 0.021 | 0.452 | 0.648 | 0.293 |
| Central Afr. Rep. | Chlorocebus sabaeus | $1.7 \times 10^5$ | 0.009 | 0.018 | 0.515 | 0.535 | 0.275 |
| Ethiopia | Chlorocebus sabaeus | $1.4 \times 10^5$ | 0.009 | 0.021 | 0.444 | 0.552 | 0.245 |
| Gambia | Chlorocebus sabaeus | $1.4 \times 10^5$ | 0.009 | 0.007 | 1.423 | 0.577 | 0.821 |
| Kenya | Chlorocebus sabaeus | $1.5 \times 10^5$ | 0.009 | 0.022 | 0.437 | 0.588 | 0.257 |
| Nevis | Chlorocebus sabaeus | $1 \times 10^5$ | 0.009 | 0.016 | 0.597 | 0.599 | 0.358 |
| South Africa | Chlorocebus sabaeus | $1.8 \times 10^5$ | 0.009 | 0.016 | 0.594 | 0.574 | 0.341 |
| Saint Kitts | Chlorocebus sabaeus | $1.2 \times 10^5$ | 0.009 | 0.017 | 0.563 | 0.598 | 0.336 |
| Zambia | Chlorocebus sabaeus | $1.7 \times 10^5$ | 0.009 | 0.022 | 0.427 | 0.585 | 0.250 |
| African | Homo sapiens | $5.6 \times 10^4$ | 0.010 | 0.020 | 0.484 | 0.721 | 0.349 |
| Admixed American | Homo sapiens | $4.5 \times 10^4$ | 0.010 | 0.019 | 0.500 | 0.690 | 0.345 |
| East Asian | Homo sapiens | $4 \times 10^4$ | 0.010 | 0.027 | 0.362 | 0.688 | 0.249 |
| European | Homo sapiens | $4.2 \times 10^4$ | 0.010 | 0.027 | 0.361 | 0.688 | 0.248 |
| South Asian | Homo sapiens | $4.4 \times 10^4$ | 0.010 | 0.030 | 0.324 | 0.691 | 0.224 |

- $N_e$ is the estimated effective population size.
- $\mathbb{P}[\mathcal{B}_0]$ (eq. 5) is the probability for a new mutation to be a non-adaptive beneficial mutation. These mutations have a selection coefficient predicted at the phylogenetic-scale larger than 1, thus toward a more fit amino-acid.
- $\mathbb{P}[\mathcal{B}]$ (eq. 15) is the probability for a mutation to be beneficial. These mutations have a selection coefficient at the population-scale larger than 1.
- $\mathbb{P}[\mathcal{B} \mid \mathcal{B}_0]$ (eq. 12) is the probability for a mutation to be beneficial at the population scale, given that it is predicted to be a non-adaptive beneficial mutation at the phylogenetic scale (the *precision*).
- $\mathbb{P}[\mathcal{B}_0 \mid \mathcal{B}]$ (eq. 14) is the probability for a mutation to be non-adaptive given that it is beneficial at the population scale (the *recall*). This probability is obtained using Bayes' formula.

#### 6.2 Misannotations of ancestral alleles

A potential caveat of our method to is that if a deleterious derived allele segregates at low frequency but is wrongly annotated as ancestral, it will result in a SNP that will be both predicted as advantageous and segregating at high frequency, wrongly suggesting that this variant is correctly inferred as beneficial. In this context misannotations of ancestral alleles could inflate our estimation of the proportion of beneficial non-adaptive mutations. However, we show that this problem impacts our estimation in two ways, albeit, only slightly.

First, if misannotation is influencing the recall for $\mathcal{B}_0$ ($\mathbb{P}[\mathcal{B}_0 \mid \mathcal{B}]$), then the recall should increase as a function of divergence to the sister species ($d_S$: number of synonymous substitutions per sites) since divergence would increase the rate of misannotations. Shown below is recall $\mathbb{P}[\mathcal{B}_0 \mid \mathcal{B}]$ as a function of divergence ($d_S$) in which the latter is not an explanatory variable.

Figure S9: $\mathbb{P}[\mathcal{B}_0 \mid \mathcal{B}]$ as a function of distance to the sister species.

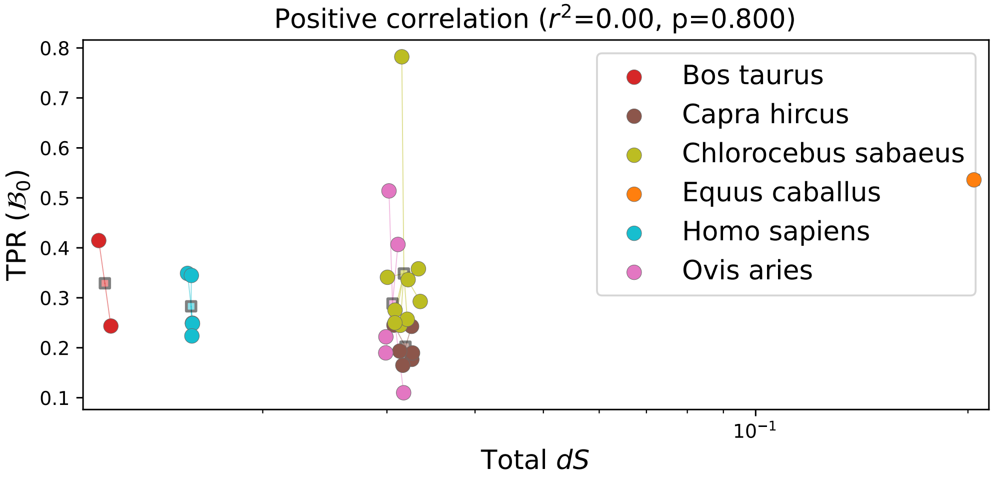

Second, the distribution of selection coefficients ($S_0$) that we observed for segregating mutations and for divergence also suggests that we effectively controlled for misannotations. Under a scenario where beneficial non-adaptive mutations would only be the result of misannotations, one would expect that the positive part of the fitness effects (i.e. $\mathbb{P}[S_0]$) of observed polymorphism currently segregating in the population should reflect the negative one (i.e. $\mathbb{P}[-S_0]$). Indeed, the number of misannotated beneficial mutations should be proportional to the number of deleterious mutations with the same selection coefficient. The ratio between advantageous and deleterious polymorphisms of the same selection coefficient should therefore be a constant that represents the rate of misannotation (typically influenced by the distance to the sister species). However, this is not what we observe in the DFE of polymorphisms (shown below in the African population of *Homo sapiens*) thus giving us the confidence that misannotations are not common.

Figure S10: Ratio of $\mathbb{P}[S_0]$ over $\mathbb{P}[-S_0]$ for polymorphisms of the African population of *Homo sapiens*.

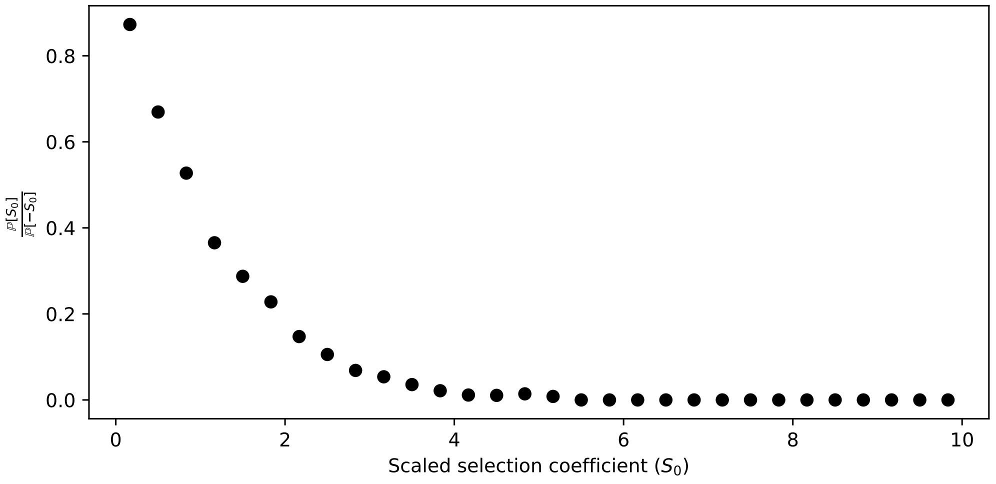

Overall, the observed DFE of substitution is much better explained by the expected DFE of new mutations (section S8.2 than by mispolarizations).

### 7 Excluding genes under adaptation

#### 7.1 Probability for beneficial mutations to be non-adaptive

Table S8: $\mathbb{P}[\mathcal{B}_0 \mid \mathcal{B}]$ excluding or not genes under adaptation.

| Population | Species | Control | Case |
|---|---|---|---|
| Equus c. | Equus caballus | 0.536 | 0.880 |
| Iran | Bos taurus | 0.243 | 0.249 |
| Uganda | Bos taurus | 0.415 | 0.429 |
| Australia | Capra hircus | 0.177 | 0.190 |
| France | Capra hircus | 0.190 | 0.201 |
| Iran (C. aegagrus) | Capra hircus | 0.165 | 0.169 |
| Iran | Capra hircus | 0.193 | 0.176 |
| Italy | Capra hircus | 0.243 | 0.261 |
| Morocco | Capra hircus | 0.245 | 0.283 |
| Iran | Ovis aries | 0.407 | 0.454 |
| Iran (O. orientalis) | Ovis aries | 0.190 | 0.207 |
| Iran (O. vignei) | Ovis aries | 0.110 | 0.135 |
| Various | Ovis aries | 0.222 | 0.246 |
| Morocco | Ovis aries | 0.514 | 0.627 |
| Barbados | Chlorocebus sabaeus | 0.293 | 0.325 |
| Central Afr. Rep. | Chlorocebus sabaeus | 0.275 | 0.273 |
| Ethiopia | Chlorocebus sabaeus | 0.245 | 0.242 |
| Gambia | Chlorocebus sabaeus | 0.821 | 0.873 |
| Kenya | Chlorocebus sabaeus | 0.257 | 0.254 |
| Nevis | Chlorocebus sabaeus | 0.358 | 0.395 |
| South Africa | Chlorocebus sabaeus | 0.341 | 0.375 |
| Saint Kitts | Chlorocebus sabaeus | 0.336 | 0.355 |
| Zambia | Chlorocebus sabaeus | 0.250 | 0.263 |
| African | Homo sapiens | 0.349 | 0.363 |
| Admixed American | Homo sapiens | 0.345 | 0.312 |
| East Asian | Homo sapiens | 0.249 | 0.233 |
| European | Homo sapiens | 0.248 | 0.211 |
| South Asian | Homo sapiens | 0.224 | 0.218 |

Genes under adaptation retrieved from Latrille *et al.* [9]. Comparison of $\mathbb{P}[\mathcal{B}_0 \mid \mathcal{B}]$ for the whole genome (control) and when excluding genes under adaptation (case). The non-parametric Wilcoxon signed-rank test evaluates the null hypothesis that the distribution of the differences (case versus control) is symmetric about zero. Applied to the paired samples (case and control), Wilcoxon signed-rank test results in $s = 80$ with $p$-value $= 0.002$, for the one-sided test (case higher than control). The proportion of non-adaptive beneficial mutations ($\mathbb{P}[\mathcal{B}_0 \mid \mathcal{B}]$) is higher when excluding genes under adaptation, consistent with our expectation that genes with uniformly conserved functions should fit better the nearly-neutral equilibrium model.

### 8 Contrasting selection at the phylogenetic and population-genetic scale

#### 8.1 Selection at the population-genetic scale

The proportion of deleterious ($\mathbb{P}[\mathcal{D}]$), nearly-neutral ($\mathbb{P}[\mathcal{N}]$) and of beneficial ($\mathbb{P}[\mathcal{B}]$) mutations estimated at the population-genetic scale across the different populations is shown for each class of selection ($x \in \{\mathcal{D}_0, \mathcal{N}_0, \mathcal{B}_0\}$).

Figure S11: Estimation of selection at the population scale for $\mathcal{D}_0$ mutations.

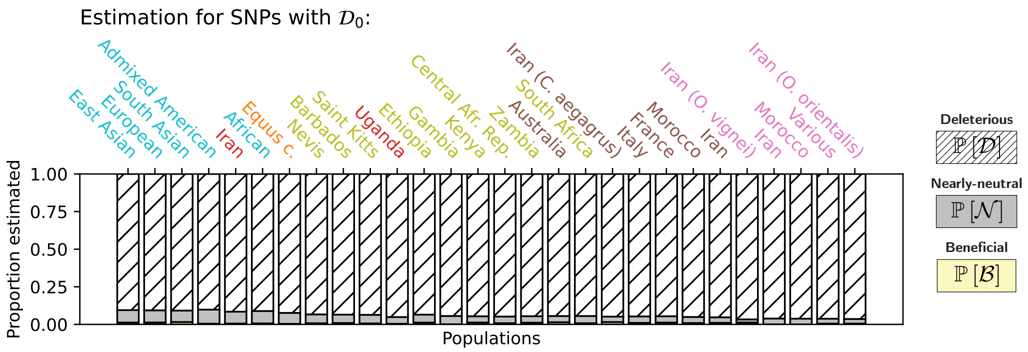

Figure S12: Estimation of selection at the population scale for $\mathcal{N}_0$ mutations.

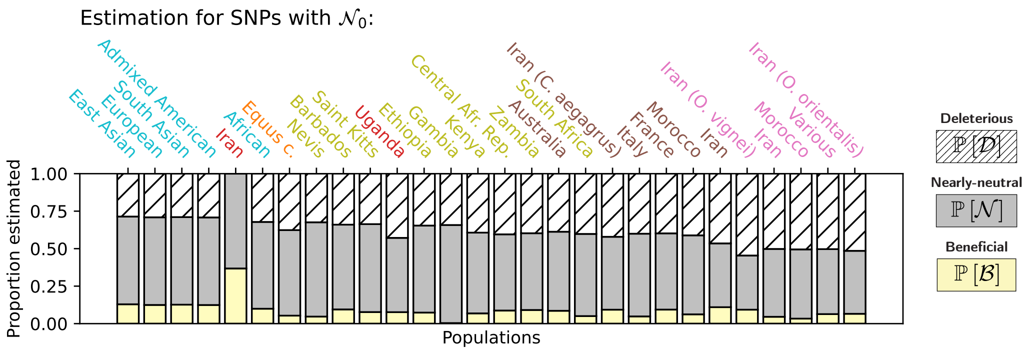

Figure S13: Estimation of selection at the population scale for $\mathcal{B}_0$ mutations.

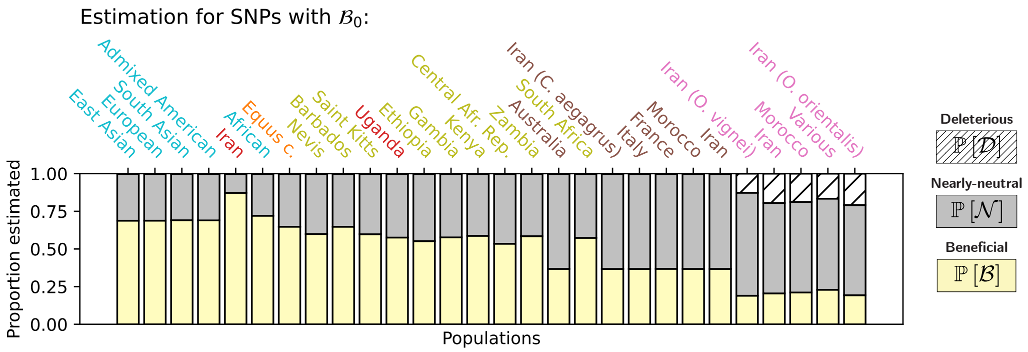

#### 8.2 Distribution of scaled selection coefficients ($S_0$)

Theoretically, the selection coefficient ($S_0$) for any mutation is related to its probability of fixation by the equation $\mathbb{P}_{\text{fix}}(S_0) = S_0/(1-\exp(-S_0))$. It is thus possible to predict the distribution of $S_0$ of substitutions given the distribution of $S_0$ for mutations (e.g. panel A below) by weighting the different mutations by their $\mathbb{P}_{\text{fix}}(S_0)$. The predicted DFE of substitutions (panel B below) matches the observed DFE of substitutions (panel C below) in the terminal lineage, meaning that the $S_0$ obtained by phylogenetic mutation-selection codon models is a strong predictor of selection effectively exerted on mutations in terminal lineages.

The figures shown below are obtained in the Saint Kitts population of *Chlorocebus sabaeus*. These figures across the 28 populations are available at Zenodo (https://doi.org/10.5281/zenodo.7878953) in file *distribution_S0_28_populations.pdf*.

Figure S14: Distribution of scaled selection coefficients ($S_0$).

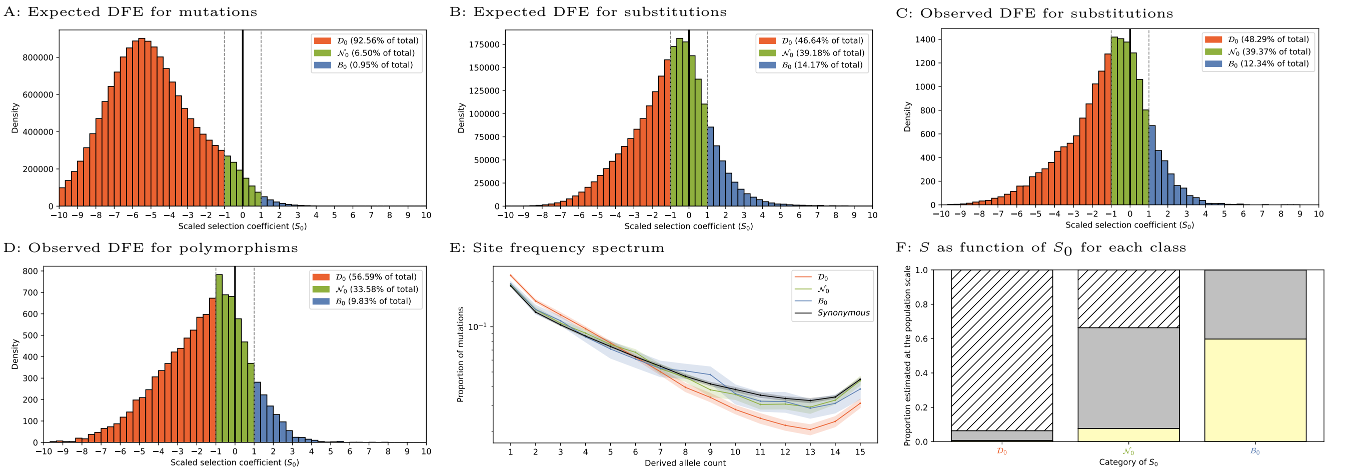

- A: Distribution of scaled selection coefficients ($S_0$), predicted for all possible non-synonymous DNA mutations away from the ancestral exome. Mutations are divided into three classes of selection: deleterious ($\mathcal{D}_0$), nearly-neutral ($\mathcal{N}_0$) and beneficial non-adaptive ($\mathcal{B}_0$).
- B: Expected distribution of $S_0$ for substitutions obtained by transforming the distribution of $S_0$ for mutations (panel A) by weighting each mutation by its scaled probability of fixation given by $\mathbb{P}_{\text{fix}}(S_0) = S_0/(1 - \exp(-S_0))$.
- C: Distribution of scaled selection coefficients ($S_0$) for all observed substitutions along the terminal branch of the phylogenetic tree.
- D: Distribution of scaled selection coefficients ($S_0$) for all observed polymorphisms currently segregating in the population.
- E: The site-frequency spectrum (SFS) in the population for a random sample of 16 alleles (means in solid lines and standard deviations in color shades) for each class of selection and for synonymous mutations, supposedly neutral (black). The SFS represents the proportion of mutations (y-axis) with a given number of derived alleles in the population (x-axis). At high frequencies, deleterious mutations are underrepresented.
- F: Proportion of beneficial $\mathbb{P}[\mathcal{D}]$, nearly-neutral $\mathbb{P}[\mathcal{N}]$, and deleterious mutations $\mathbb{P}[\mathcal{B}]$ estimated at the population scale for each class of selection at the phylogenetic scale. Proportions depicted here are not weighted by their mutational opportunities.

### 9 Filtering of variable sites

#### 9.1 Masking CpG transitions

We also tested our prediction while masking CpG transitions since the mutation rate of CpG dinucleotides is higher than the mutation rate of other dinucleotides, which can affect the estimation of the DFE. Any nucleotide transition including a CpG dinucleotide either in the ancestral or derived state is considered as a CpG transition. We masked all CpG transitions in the mutational opportunities, the substitutions in the terminal branches, and the polymorphisms segregating in the population. We then re-estimated the DFE and the proportion of non-adaptive beneficial mutations.

#### 9.2 Probability for beneficial mutations to be non-adaptive

Table S9: $\mathbb{P}[\mathcal{B}_0 \mid \mathcal{B}]$ while masking CpG transitions.

| Population | Species | $N_e$ | $\mathbb{P}[\mathcal{B}_0]$ | $\mathbb{P}[\mathcal{B}]$ | $\frac{\mathbb{P}[\mathcal{B}_0]}{\mathbb{P}[\mathcal{B}]}$ | $\mathbb{P}[\mathcal{B} \mid \mathcal{B}_0]$ | $\mathbb{P}[\mathcal{B}_0 \mid \mathcal{B}]$ |
|---|---|---|---|---|---|---|---|
| Equus c. | Equus caballus | $7.5 \times 10^4$ | 0.012 | 0.028 | 0.421 | 0.543 | 0.229 |
| Iran | Bos taurus | $5.6 \times 10^4$ | 0.011 | 0.038 | 0.283 | 0.797 | 0.226 |
| Uganda | Bos taurus | $1.3 \times 10^5$ | 0.011 | 0.022 | 0.479 | 0.406 | 0.195 |
| Australia | Capra hircus | $1.7 \times 10^5$ | 0.011 | 0.024 | 0.454 | 0.158 | 0.072 |
| France | Capra hircus | $1.9 \times 10^5$ | 0.011 | 0.034 | 0.320 | 0.258 | 0.082 |
| Iran (C. aegagrus) | Capra hircus | $1.9 \times 10^5$ | 0.011 | 0.032 | 0.341 | 0.268 | 0.091 |
| Iran | Capra hircus | $2.3 \times 10^5$ | 0.011 | 0.026 | 0.417 | 0.294 | 0.123 |
| Italy | Capra hircus | $1.9 \times 10^5$ | 0.011 | 0.023 | 0.472 | 0.100 | 0.047 |
| Morocco | Capra hircus | $2.2 \times 10^5$ | 0.011 | 0.018 | 0.593 | 0.141 | 0.083 |
| Iran | Ovis aries | $3.8 \times 10^5$ | 0.011 | 0.012 | 0.981 | 0.243 | 0.239 |
| Iran (O. orientalis) | Ovis aries | $4.5 \times 10^5$ | 0.011 | 0.019 | 0.603 | 0.155 | 0.093 |
| Iran (O. vignei) | Ovis aries | $3.7 \times 10^5$ | 0.011 | 0.025 | 0.445 | 0.252 | 0.112 |
| Various | Ovis aries | $4.1 \times 10^5$ | 0.011 | 0.016 | 0.704 | 0.198 | 0.139 |
| Morocco | Ovis aries | $4 \times 10^5$ | 0.011 | 0.008 | 1.389 | 0.165 | 0.229 |
| Barbados | Chlorocebus sabaeus | $1.1 \times 10^5$ | 0.009 | 0.013 | 0.752 | 0.231 | 0.174 |
| Central Afr. Rep. | Chlorocebus sabaeus | $1.7 \times 10^5$ | 0.009 | 0.018 | 0.519 | 0.209 | 0.108 |
| Ethiopia | Chlorocebus sabaeus | $1.4 \times 10^5$ | 0.009 | 0.025 | 0.385 | 0.241 | 0.093 |
| Gambia | Chlorocebus sabaeus | $1.4 \times 10^5$ | 0.009 | 0.007 | 1.346 | 0.204 | 0.275 |
| Kenya | Chlorocebus sabaeus | $1.5 \times 10^5$ | 0.009 | 0.021 | 0.443 | 0.239 | 0.106 |
| Nevis | Chlorocebus sabaeus | $1 \times 10^5$ | 0.009 | 0.014 | 0.690 | 0.146 | 0.101 |
| South Africa | Chlorocebus sabaeus | $1.8 \times 10^5$ | 0.009 | 0.019 | 0.498 | 0.272 | 0.135 |
| Saint Kitts | Chlorocebus sabaeus | $1.2 \times 10^5$ | 0.009 | 0.017 | 0.567 | 0.217 | 0.123 |
| Zambia | Chlorocebus sabaeus | $1.7 \times 10^5$ | 0.009 | 0.024 | 0.388 | 0.308 | 0.119 |
| African | Homo sapiens | $5.6 \times 10^4$ | 0.010 | 0.026 | 0.367 | 0.411 | 0.151 |
| Admixed American | Homo sapiens | $4.5 \times 10^4$ | 0.010 | 0.017 | 0.551 | 0.374 | 0.206 |
| East Asian | Homo sapiens | $4 \times 10^4$ | 0.010 | 0.029 | 0.332 | 0.370 | 0.123 |
| European | Homo sapiens | $4.2 \times 10^4$ | 0.010 | 0.031 | 0.304 | 0.371 | 0.113 |
| South Asian | Homo sapiens | $4.4 \times 10^4$ | 0.010 | 0.032 | 0.302 | 0.369 | 0.111 |

- $N_e$ is the estimated effective population size.
- $\mathbb{P}[\mathcal{B}_0]$ (eq. 5) is the probability for a new mutation a non-adaptive beneficial mutation. These mutations have a selection coefficient predicted at the phylogenetic-scale larger than 1, thus toward a more fit amino-acid.
- $\mathbb{P}[\mathcal{B}]$ (eq. 15) is the probability for a mutation to be beneficial. These mutations have a selection coefficient at the population-scale larger than 1.
- $\mathbb{P}[\mathcal{B} \mid \mathcal{B}_0]$ (eq. 12) is the probability for a mutation to be beneficial at the population scale, given it is predicted to be a non-adaptive beneficial mutation at the phylogenetic scale (the *precision*).
- $\mathbb{P}[\mathcal{B}_0 \mid \mathcal{B}]$ (eq. 14) is the probability for a mutation to be non-adaptive given it is beneficial at the population scale (the *recall*). This probability is obtained using Bayes' formula.

#### 9.3 Precision and recall

Table S10: Precision and recall while masking CpG transitions.

| | | | Deleterious mutations $\mathcal{D} := S < -1$ $\mathcal{D}_0 := S_0 < -1$ | | Nearly-neutral mutations $\mathcal{N} := -1 < S < 1$ $\mathcal{N}_0 := -1 < S_0 < 1$ | | Beneficial mutations $\mathcal{B} := S > 1$ $\mathcal{B}_0 := S_0 > 1$ | |
|---|---|---|---|---|---|---|---|---|
| **Population** | **Species** | $N_e$ | **Precision** $\mathbb{P}[\mathcal{D} \mid \mathcal{D}_0]$ | **Recall** $\mathbb{P}[\mathcal{D}_0 \mid \mathcal{D}]$ | **Precision** $\mathbb{P}[\mathcal{N} \mid \mathcal{N}_0]$ | **Recall** $\mathbb{P}[\mathcal{N}_0 \mid \mathcal{N}]$ | **Precision** $\mathbb{P}[\mathcal{B} \mid \mathcal{B}_0]$ | **Recall** $\mathbb{P}[\mathcal{B}_0 \mid \mathcal{B}]$ |
| Equus c. | Equus caballus | $7.5 \times 10^4$ | 0.887 | 0.984 | 0.740 | 0.356 | 0.543 | 0.229 |
| Iran | Bos taurus | $5.6 \times 10^4$ | 0.888 | 1.000 | 0.632 | 0.302 | 0.797 | 0.226 |
| Uganda | Bos taurus | $1.3 \times 10^5$ | 0.929 | 0.973 | 0.501 | 0.349 | 0.406 | 0.195 |
| Australia | Capra hircus | $1.7 \times 10^5$ | 0.926 | 0.972 | 0.609 | 0.421 | 0.158 | 0.072 |
| France | Capra hircus | $1.9 \times 10^5$ | 0.927 | 0.959 | 0.354 | 0.321 | 0.258 | 0.082 |
| Iran (C. aegagrus) | Capra hircus | $1.9 \times 10^5$ | 0.931 | 0.967 | 0.502 | 0.423 | 0.268 | 0.091 |
| Iran | Capra hircus | $2.3 \times 10^5$ | 0.937 | 0.966 | 0.529 | 0.451 | 0.294 | 0.123 |
| Italy | Capra hircus | $1.9 \times 10^5$ | 0.927 | 0.970 | 0.550 | 0.390 | 0.100 | 0.047 |
| Morocco | Capra hircus | $2.2 \times 10^5$ | 0.933 | 0.965 | 0.534 | 0.401 | 0.141 | 0.083 |
| Iran | Ovis aries | $3.8 \times 10^5$ | 0.947 | 0.957 | 0.446 | 0.393 | 0.243 | 0.239 |
| Iran (O. orientalis) | Ovis aries | $4.5 \times 10^5$ | 0.951 | 0.961 | 0.396 | 0.386 | 0.155 | 0.093 |
| Iran (O. vignei) | Ovis aries | $3.7 \times 10^5$ | 0.954 | 0.960 | 0.379 | 0.432 | 0.252 | 0.112 |
| Various | Ovis aries | $4.1 \times 10^5$ | 0.949 | 0.956 | 0.434 | 0.418 | 0.198 | 0.139 |
| Morocco | Ovis aries | $4 \times 10^5$ | 0.946 | 0.959 | 0.453 | 0.374 | 0.165 | 0.229 |
| Barbados | Chlorocebus sabaeus | $1.1 \times 10^5$ | 0.916 | 0.971 | 0.523 | 0.306 | 0.231 | 0.174 |
| Central Afr. Rep. | Chlorocebus sabaeus | $1.7 \times 10^5$ | 0.933 | 0.969 | 0.486 | 0.353 | 0.209 | 0.108 |
| Ethiopia | Chlorocebus sabaeus | $1.4 \times 10^5$ | 0.918 | 0.973 | 0.533 | 0.348 | 0.241 | 0.093 |
| Gambia | Chlorocebus sabaeus | $1.4 \times 10^5$ | 0.925 | 0.971 | 0.572 | 0.343 | 0.204 | 0.275 |
| Kenya | Chlorocebus sabaeus | $1.5 \times 10^5$ | 0.931 | 0.971 | 0.521 | 0.378 | 0.239 | 0.106 |
| Nevis | Chlorocebus sabaeus | $1 \times 10^5$ | 0.916 | 0.972 | 0.535 | 0.313 | 0.146 | 0.101 |
| South Africa | Chlorocebus sabaeus | $1.8 \times 10^5$ | 0.928 | 0.970 | 0.546 | 0.378 | 0.272 | 0.135 |
| Saint Kitts | Chlorocebus sabaeus | $1.2 \times 10^5$ | 0.917 | 0.971 | 0.507 | 0.309 | 0.217 | 0.123 |
| Zambia | Chlorocebus sabaeus | $1.7 \times 10^5$ | 0.930 | 0.969 | 0.495 | 0.372 | 0.308 | 0.119 |
| African | Homo sapiens | $5.6 \times 10^4$ | 0.888 | 0.978 | 0.587 | 0.296 | 0.411 | 0.151 |
| Admixed American | Homo sapiens | $4.5 \times 10^4$ | 0.874 | 0.979 | 0.591 | 0.256 | 0.374 | 0.206 |
| East Asian | Homo sapiens | $4 \times 10^4$ | 0.877 | 0.980 | 0.593 | 0.280 | 0.370 | 0.123 |
| European | Homo sapiens | $4.2 \times 10^4$ | 0.885 | 0.980 | 0.591 | 0.301 | 0.371 | 0.113 |
| South Asian | Homo sapiens | $4.4 \times 10^4$ | 0.886 | 0.979 | 0.591 | 0.303 | 0.369 | 0.111 |

- $N_e$ is the estimated effective population size.
- *Precision* is the estimation of the selection coefficient at population scale ($S$) given that $S_0$ is known.
- *Recall* is the estimation of $S_0$ given selection coefficient at the population scale ($S$) is known.
- *Recall* for beneficial mutations ($\mathbb{P}[\mathcal{B}_0 \mid \mathcal{B}]$) is the probability for a mutation to be non-adaptive given it is beneficial at the population scale.

Altogether, comparing this table other estimations, we acknowledge that the exact proportion of non-adaptive among beneficial mutations is dependent on the whether we masked CpG sites. However, we can still see that non-adaptive beneficial mutations are positively selected compared to neutral and deleterious ones, and this result is not dependent on whether we masked CpG sites or not.

#### 9.4 Example site-frequency spectrum

The site-frequency spectrum (SFS, panel A) and the congruency between fitness effects estimates ($S_0$ versus $S_0$, panel B) shown below are obtained in the African population of *Homo sapiens* and the South African population of *Chlorocebus sabaeus*.

Figure S15: Site-frequency spectrum in African population of *Homo sapiens*, masking CpG transitions.

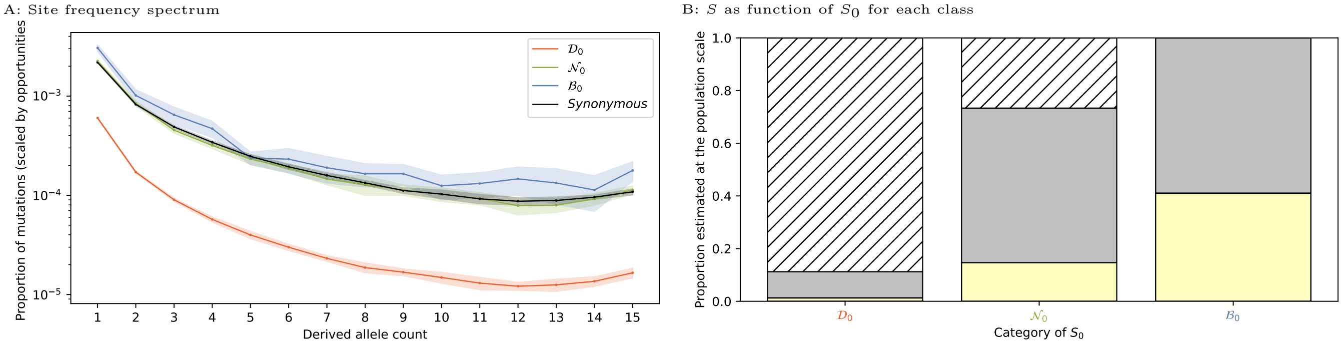

Figure S16: Site-frequency spectrum in South African population of *Chlorocebus sabaeus*, masking CpG transitions.

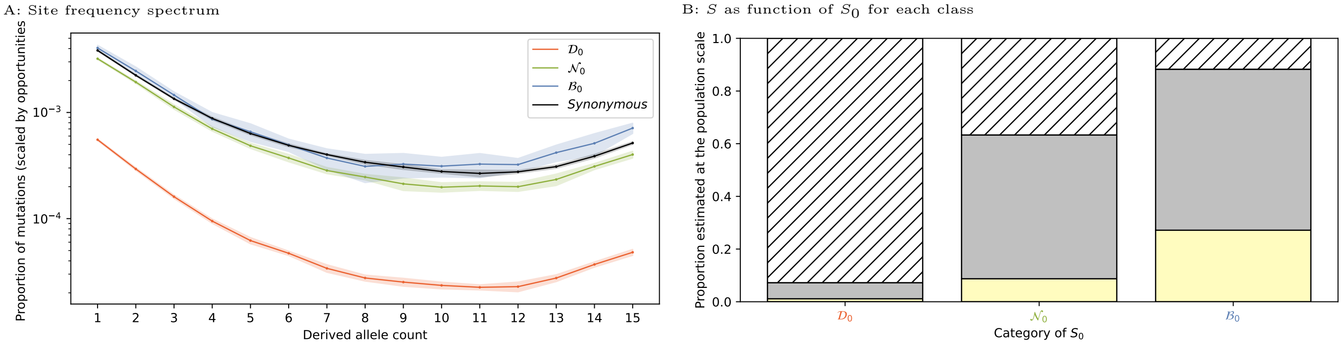

- A: The site-frequency spectrum (SFS) in the population for a random sample of 16 alleles (means in solid lines and standard deviations in color shades) for each class of selection and for synonymous mutations, supposedly neutral (black). The SFS represents the proportion of mutations (y-axis) with a given number of derived alleles in the population (x-axis). Here SFS are weighted by the mutational opportunities in each class of selection (synonymous, $\mathcal{D}_0$, $\mathcal{N}_0$ and $\mathcal{B}_0$).
- B: Proportion of beneficial $\mathbb{P}[\mathcal{D}]$, nearly-neutral $\mathbb{P}[\mathcal{N}]$, and deleterious mutations $\mathbb{P}[\mathcal{B}]$ estimated at the population scale for each class of selection at the phylogenetic scale. Proportions depicted here are not weighted by their mutational opportunities.

### 10 Including divergence data for the estimation of $S$

#### 10.1 Including divergence in polyDFE

Divergence data (number of substitutions per site) can also be included into polyDFE to estimate the DFE. For the class of a given class of selection coefficient ($x \in \{\mathcal{D}_0, \mathcal{N}_0, \mathcal{B}_0\}$), the number of substitutions has already been computed and is given by $D(x)$ (see eq. 6), and the number of sites is given by $L(x)$ (see eq. 4). Otherwise, the procedure is the same as described section *Scaled selection coefficients (S) in a population-based method.*

#### 10.2 Probability for beneficial mutations to be non-adaptive

Table S11: $\mathbb{P}[\mathcal{B}_0 \mid \mathcal{B}]$ when including divergence to estimate $S$.

| Population | Species | $N_e$ | $\mathbb{P}[\mathcal{B}_0]$ | $\mathbb{P}[\mathcal{B}]$ | $\frac{\mathbb{P}[\mathcal{B}_0]}{\mathbb{P}[\mathcal{B}]}$ | $\mathbb{P}[\mathcal{B} \mid \mathcal{B}_0]$ | $\mathbb{P}[\mathcal{B}_0 \mid \mathcal{B}]$ |
|---|---|---|---|---|---|---|---|
| Equus c. | Equus caballus | $7.5 \times 10^4$ | 0.012 | 0.057 | 0.213 | 0.498 | 0.106 |
| Iran | Bos taurus | $5.6 \times 10^4$ | 0.011 | 0.006 | 1.860 | 0.368 | 0.684 |
| Uganda | Bos taurus | $1.3 \times 10^5$ | 0.011 | 0.012 | 0.900 | 0.307 | 0.276 |
| Australia | Capra hircus | $1.7 \times 10^5$ | 0.011 | 0.007 | 1.651 | 0.178 | 0.294 |
| France | Capra hircus | $1.9 \times 10^5$ | 0.011 | 0.007 | 1.614 | 0.183 | 0.295 |
| Iran (C. aegagrus) | Capra hircus | $1.9 \times 10^5$ | 0.011 | 0.012 | 0.923 | 0.177 | 0.163 |
| Iran | Capra hircus | $2.3 \times 10^5$ | 0.011 | 0.008 | 1.483 | 0.184 | 0.273 |
| Italy | Capra hircus | $1.9 \times 10^5$ | 0.011 | 0.007 | 1.686 | 0.178 | 0.299 |
| Morocco | Capra hircus | $2.2 \times 10^5$ | 0.011 | 0.006 | 1.892 | 0.184 | 0.349 |
| Iran | Ovis aries | $3.8 \times 10^5$ | 0.012 | 0.010 | 1.132 | 0.189 | 0.214 |
| Iran (O. orientalis) | Ovis aries | $4.5 \times 10^5$ | 0.011 | 0.011 | 1.012 | 0.213 | 0.216 |
| Iran (O. vignei) | Ovis aries | $3.7 \times 10^5$ | 0.012 | 0.021 | 0.561 | 0.186 | 0.105 |
| Various | Ovis aries | $4.1 \times 10^5$ | 0.011 | 0.011 | 1.069 | 0.215 | 0.229 |
| Morocco | Ovis aries | $4 \times 10^5$ | 0.012 | 0.012 | 0.973 | 0.217 | 0.211 |
| Barbados | Chlorocebus sabaeus | $1.1 \times 10^5$ | 0.009 | 0.004 | 2.120 | 0.341 | 0.723 |
| Central Afr. Rep. | Chlorocebus sabaeus | $1.7 \times 10^5$ | 0.009 | 0.013 | 0.720 | 0.368 | 0.265 |
| Ethiopia | Chlorocebus sabaeus | $1.4 \times 10^5$ | 0.009 | 0.007 | 1.352 | 0.368 | 0.498 |
| Gambia | Chlorocebus sabaeus | $1.4 \times 10^5$ | 0.009 | 0.010 | 0.956 | 0.368 | 0.352 |
| Kenya | Chlorocebus sabaeus | $1.5 \times 10^5$ | 0.009 | 0.013 | 0.727 | 0.368 | 0.268 |
| Nevis | Chlorocebus sabaeus | $1 \times 10^5$ | 0.009 | 0.004 | 2.133 | 0.368 | 0.784 |
| South Africa | Chlorocebus sabaeus | $1.8 \times 10^5$ | 0.009 | 0.011 | 0.874 | 0.368 | 0.322 |
| Saint Kitts | Chlorocebus sabaeus | $1.2 \times 10^5$ | 0.009 | 0.005 | 1.905 | 0.368 | 0.701 |
| Zambia | Chlorocebus sabaeus | $1.7 \times 10^5$ | 0.009 | 0.014 | 0.698 | 0.368 | 0.257 |
| African | Homo sapiens | $5.6 \times 10^4$ | 0.010 | 0.013 | 0.726 | 0.437 | 0.317 |
| Admixed American | Homo sapiens | $4.5 \times 10^4$ | 0.010 | 0.011 | 0.886 | 0.429 | 0.380 |
| East Asian | Homo sapiens | $4 \times 10^4$ | 0.010 | 0.008 | 1.213 | 0.426 | 0.517 |
| European | Homo sapiens | $4.1 \times 10^4$ | 0.010 | 0.009 | 1.104 | 0.422 | 0.466 |
| South Asian | Homo sapiens | $4.4 \times 10^4$ | 0.010 | 0.009 | 1.016 | 0.426 | 0.433 |

- $N_e$ is the estimated effective population size.
- $\mathbb{P}[\mathcal{B}_0]$ (eq. 5) is the probability for a new mutation a non-adaptive mutation. These mutations have a selection coefficient predicted at the phylogenetic-scale larger than 1, thus toward a more fit amino-acid.
- $\mathbb{P}[\mathcal{B}]$ (eq. 15) is the probability for a mutation to be beneficial. These mutations have a selection coefficient at the population-scale larger than 1.
- $\mathbb{P}[\mathcal{B} \mid \mathcal{B}_0]$ (eq. 12) is the probability for a mutation to be beneficial at the population scale, given it is predicted to be a non-adaptive mutation at the phylogenetic scale (the *precision*).
- $\mathbb{P}[\mathcal{B}_0 \mid \mathcal{B}]$ (eq. 14) is the probability for a mutation to be non-adaptive given it is beneficial at the population scale (the *recall*). This probability is obtained using Bayes' formula.

#### 10.3 Precision and recall

Table S12: Precision and recall while including divergence to estimate $S$.

| | | | Deleterious mutations $\mathcal{D} \coloneqq S < -1$ $\mathcal{D}_0 \coloneqq S_0 < -1$ | | Nearly-neutral mutations $\mathcal{N} \coloneqq -1 < S < 1$ $\mathcal{N}_0 \coloneqq -1 < S_0 < 1$ | | Beneficial mutations $\mathcal{B} \coloneqq S > 1$ $\mathcal{B}_0 \coloneqq S_0 > 1$ | |
|---|---|---|---|---|---|---|---|---|
| **Population** | **Species** | $N_e$ | **Precision** $\mathbb{P}[\mathcal{D} \mid \mathcal{D}_0]$ | **Recall** $\mathbb{P}[\mathcal{D}_0 \mid \mathcal{D}]$ | **Precision** $\mathbb{P}[\mathcal{N} \mid \mathcal{N}_0]$ | **Recall** $\mathbb{P}[\mathcal{N}_0 \mid \mathcal{N}]$ | **Precision** $\mathbb{P}[\mathcal{B} \mid \mathcal{B}_0]$ | **Recall** $\mathbb{P}[\mathcal{B}_0 \mid \mathcal{B}]$ |
| Equus c. | Equus caballus | $7.5 \times 10^4$ | 0.937 | 0.955 | 0.111 | 0.192 | 0.498 | 0.106 |
| Iran | Bos taurus | $5.6 \times 10^4$ | 0.935 | 0.970 | 0.565 | 0.356 | 0.368 | 0.684 |
| Uganda | Bos taurus | $1.3 \times 10^5$ | 0.951 | 0.967 | 0.516 | 0.422 | 0.307 | 0.276 |
| Australia | Capra hircus | $1.7 \times 10^5$ | 0.952 | 0.970 | 0.598 | 0.452 | 0.178 | 0.294 |
| France | Capra hircus | $1.9 \times 10^5$ | 0.951 | 0.969 | 0.574 | 0.433 | 0.183 | 0.295 |
| Iran (C. aegagrus) | Capra hircus | $1.9 \times 10^5$ | 0.957 | 0.966 | 0.515 | 0.460 | 0.177 | 0.163 |
| Iran | Capra hircus | $2.3 \times 10^5$ | 0.951 | 0.967 | 0.542 | 0.422 | 0.184 | 0.273 |
| Italy | Capra hircus | $1.9 \times 10^5$ | 0.951 | 0.969 | 0.582 | 0.436 | 0.178 | 0.299 |
| Morocco | Capra hircus | $2.2 \times 10^5$ | 0.949 | 0.968 | 0.556 | 0.412 | 0.184 | 0.349 |
| Iran | Ovis aries | $3.8 \times 10^5$ | 0.963 | 0.962 | 0.421 | 0.424 | 0.189 | 0.214 |
| Iran (O. orientalis) | Ovis aries | $4.5 \times 10^5$ | 0.968 | 0.961 | 0.413 | 0.460 | 0.213 | 0.216 |
| Iran (O. vignei) | Ovis aries | $3.7 \times 10^5$ | 0.972 | 0.959 | 0.360 | 0.528 | 0.186 | 0.105 |
| Various | Ovis aries | $4.1 \times 10^5$ | 0.966 | 0.961 | 0.430 | 0.454 | 0.215 | 0.229 |
| Morocco | Ovis aries | $4 \times 10^5$ | 0.966 | 0.958 | 0.372 | 0.416 | 0.217 | 0.211 |
| Barbados | Chlorocebus sabaeus | $1.1 \times 10^5$ | 0.940 | 0.976 | 0.654 | 0.409 | 0.341 | 0.723 |
| Central Afr. Rep. | Chlorocebus sabaeus | $1.7 \times 10^5$ | 0.950 | 0.971 | 0.526 | 0.421 | 0.368 | 0.265 |
| Ethiopia | Chlorocebus sabaeus | $1.4 \times 10^5$ | 0.942 | 0.974 | 0.609 | 0.402 | 0.368 | 0.498 |
| Gambia | Chlorocebus sabaeus | $1.4 \times 10^5$ | 0.950 | 0.974 | 0.611 | 0.452 | 0.368 | 0.352 |
| Kenya | Chlorocebus sabaeus | $1.5 \times 10^5$ | 0.950 | 0.972 | 0.545 | 0.434 | 0.368 | 0.268 |
| Nevis | Chlorocebus sabaeus | $1 \times 10^5$ | 0.940 | 0.977 | 0.668 | 0.412 | 0.368 | 0.784 |
| South Africa | Chlorocebus sabaeus | $1.8 \times 10^5$ | 0.947 | 0.972 | 0.550 | 0.411 | 0.368 | 0.322 |
| Saint Kitts | Chlorocebus sabaeus | $1.2 \times 10^5$ | 0.940 | 0.976 | 0.646 | 0.407 | 0.368 | 0.701 |
| Zambia | Chlorocebus sabaeus | $1.7 \times 10^5$ | 0.950 | 0.971 | 0.527 | 0.426 | 0.368 | 0.257 |
| African | Homo sapiens | $5.6 \times 10^4$ | 0.911 | 0.980 | 0.666 | 0.341 | 0.437 | 0.317 |
| Admixed American | Homo sapiens | $4.5 \times 10^4$ | 0.902 | 0.976 | 0.580 | 0.284 | 0.429 | 0.380 |
| East Asian | Homo sapiens | $4 \times 10^4$ | 0.904 | 0.984 | 0.744 | 0.341 | 0.426 | 0.517 |
| European | Homo sapiens | $4.1 \times 10^4$ | 0.906 | 0.984 | 0.737 | 0.344 | 0.422 | 0.466 |
| South Asian | Homo sapiens | $4.4 \times 10^4$ | 0.907 | 0.984 | 0.737 | 0.349 | 0.426 | 0.433 |

- $N_e$ is the estimated effective population size.
- *Precision* is the estimation of the selection coefficient at population scale ($S$) given that $S_0$ is known.
- *Recall* is the estimation of $S_0$ given selection coefficient at the population scale ($S$) is known.
- *Recall* for beneficial mutations ($\mathbb{P}[\mathcal{B}_0 \mid \mathcal{B}]$) is the probability for a mutation to be non-adaptive given it is beneficial at the population scale.

Altogether, comparing this table to tables 1 and S3, we acknowledge that the exact proportion of non-adaptive beneficial mutations among all beneficial ones is different whether we included or not substitutions in the terminal lineage for the estimation of $S$. However, we can still see that non-adaptive beneficial mutations are positively selected compared to neutral and deleterious ones, and this result is not sensitive to inclusion of substitutions in the terminal lineage.

#### 10.4 Misannotations of ancestral alleles

If misannotation is influencing the recall for $\mathcal{B}_0$ ($\mathbb{P}[\mathcal{B}_0 \mid \mathcal{B}]$), then the recall should increase as a function of divergence to the sister species ($d_S$: number of synonymous substitutions per sites) since divergence would increase the rate of misannotations. Shown below is recall $\mathbb{P}[\mathcal{B}_0 \mid \mathcal{B}]$ as a function of divergence ($d_S$) in which the latter is not an explanatory variable.

Figure S17: $\mathbb{P}[\mathcal{B}_0 \mid \mathcal{B}]$ while including divergence to estimate $S$, as a function of distance to the sister species.

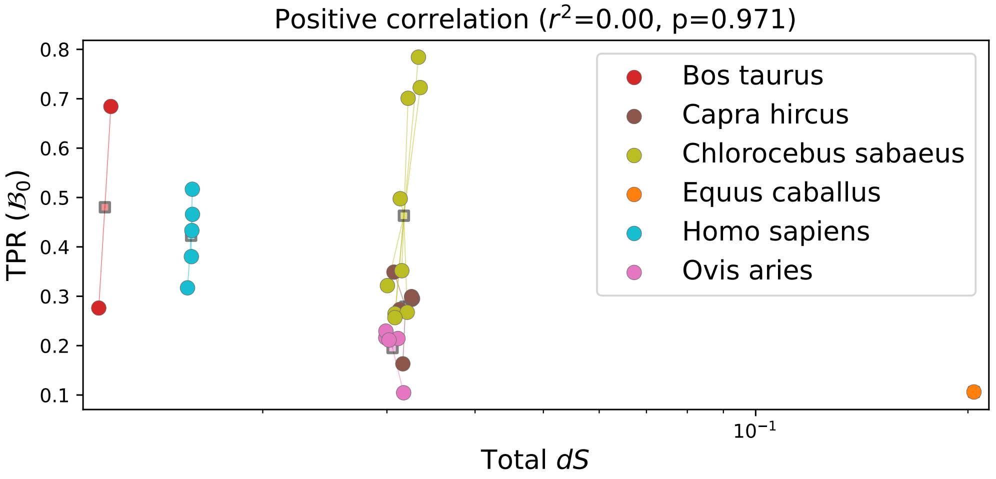

#### 10.5 Excluding genes under adaptation

Table S13: $\mathbb{P}[\mathcal{B}_0 \mid \mathcal{B}]$ excluding or not genes under adaptation, when including divergence to estimate $S$.

| Population | Species | Control | Case |
|---|---|---|---|
| Equus c. | Equus caballus | 0.106 | 0.106 |
| Iran | Bos taurus | 0.684 | 0.755 |
| Uganda | Bos taurus | 0.276 | 0.310 |
| Australia | Capra hircus | 0.294 | 0.319 |
| France | Capra hircus | 0.295 | 0.325 |
| Iran (C. aegagrus) | Capra hircus | 0.163 | 0.174 |
| Iran | Capra hircus | 0.273 | 0.299 |
| Italy | Capra hircus | 0.299 | 0.321 |
| Morocco | Capra hircus | 0.349 | 0.419 |
| Iran | Ovis aries | 0.214 | 0.241 |
| Iran (O. orientalis) | Ovis aries | 0.216 | 0.213 |
| Iran (O. vignei) | Ovis aries | 0.105 | 0.089 |
| Various | Ovis aries | 0.229 | 0.196 |
| Morocco | Ovis aries | 0.211 | 0.206 |
| Barbados | Chlorocebus sabaeus | 0.723 | 0.833 |
| Central Afr. Rep. | Chlorocebus sabaeus | 0.265 | 0.285 |
| Ethiopia | Chlorocebus sabaeus | 0.498 | 0.547 |
| Gambia | Chlorocebus sabaeus | 0.352 | 0.571 |
| Kenya | Chlorocebus sabaeus | 0.268 | 0.265 |
| Nevis | Chlorocebus sabaeus | 0.784 | 0.892 |
| South Africa | Chlorocebus sabaeus | 0.322 | 0.307 |
| Saint Kitts | Chlorocebus sabaeus | 0.701 | 0.825 |
| Zambia | Chlorocebus sabaeus | 0.257 | 0.271 |
| African | Homo sapiens | 0.317 | 0.278 |
| Admixed American | Homo sapiens | 0.380 | 0.363 |
| East Asian | Homo sapiens | 0.517 | 0.499 |
| European | Homo sapiens | 0.466 | 0.456 |
| South Asian | Homo sapiens | 0.433 | 0.304 |

Genes under adaptation retrieved from [9]. Comparison of $\mathbb{P}[\mathcal{B}_0 \mid \mathcal{B}]$ while including divergence for the whole genome (control) and when excluding genes under adaptation (case). The non-parametric Wilcoxon signed-rank test tests the null hypothesis that the distribution of the differences (case versus control) is symmetric about zero. Applied to the paired samples (case and control), Wilcoxon signed-rank test results in $s = 120$ with $p$-value $= 0.027$ for one-sided test (case higher than control). The proportion of beneficial non-adaptive mutations ($\mathbb{P}[\mathcal{B}_0 \mid \mathcal{B}]$) is higher when excluding genes under adaptation, consistent with our expectation that genes with uniformly conserved functions should fit better the nearly-neutral equilibrium model.

### 11 Discrete distribution of $S$ at the population scale

#### 11.1 polyDFE model D (discrete distribution)

Additionally to including divergence, with also tested our prediction with polyDFE model D instead of model C. In polyDFE model D, the DFE of non-synonymous mutations is given as a discrete DFE of $K$ categories (instead of a continuous distribution in model C), where the selection coefficients of each category $i$ ($1 \leq i \leq K$) are fixed parameters $S_i$, and each value $S_i$ has a probability $p_i$ (estimated), with $\sum_{i=1}^{K} p_i = 1$. We used $K = 6$ with $S_1 = -500$, $S_2 = -4$, $S_3 = -1$, $S_4 = 0$, $S_5 = 1$, $S_6 = 4$.

For each class of selection $x$, the parameters $p_i$ ($i \in \{1 \leq i \leq 6\}$) were used to compute $\mathbb{P}[\mathcal{D} \mid x]$, $\mathbb{P}[\mathcal{N} \mid x]$, and $\mathbb{P}[\mathcal{B} \mid x]$ as:

$$\mathbb{P}[\mathcal{D} \mid x] = \mathbb{P}[S < -1 \mid x] = p_1 + p_2 \tag{S.15}$$

$$\mathbb{P}[\mathcal{N} \mid x] = \mathbb{P}[-1 < S < 1 \mid x] = p_3 + p_4 + p_5, \tag{S.16}$$

$$\mathbb{P}[\mathcal{B} \mid x] = \mathbb{P}[S > 1 \mid x] = p_6. \tag{S.17}$$

#### 11.2 Probability for beneficial mutations to be non-adaptive

Table S14: $\mathbb{P}[\mathcal{B}_0 \mid \mathcal{B}]$ when using a discrete DFE to estimate $S$.

| Population | Species | $N_e$ | $\mathbb{P}[\mathcal{B}_0]$ | $\mathbb{P}[\mathcal{B}]$ | $\frac{\mathbb{P}[\mathcal{B}_0]}{\mathbb{P}[\mathcal{B}]}$ | $\mathbb{P}[\mathcal{B} \mid \mathcal{B}_0]$ | $\mathbb{P}[\mathcal{B}_0 \mid \mathcal{B}]$ |
|---|---|---|---|---|---|---|---|
| Equus c. | Equus caballus | $7.5 \times 10^4$ | 0.012 | 0.006 | 2.050 | 0.163 | 0.334 |
| Iran | Bos taurus | $5.6 \times 10^4$ | 0.011 | 0.005 | 2.125 | 0.133 | 0.283 |
| Uganda | Bos taurus | $1.3 \times 10^5$ | 0.011 | 0.010 | 1.139 | 0.140 | 0.159 |
| Australia | Capra hircus | $1.7 \times 10^5$ | 0.011 | 0.002 | 5.637 | 0.151 | 0.850 |
| France | Capra hircus | $1.9 \times 10^5$ | 0.011 | 0.002 | 6.117 | 0.154 | 0.940 |
| Iran (C. aegagrus) | Capra hircus | $1.9 \times 10^5$ | 0.011 | 0.002 | 5.842 | 0.156 | 0.911 |
| Iran | Capra hircus | $2.3 \times 10^5$ | 0.011 | 0.002 | 6.400 | 0.147 | 0.943 |
| Italy | Capra hircus | $1.9 \times 10^5$ | 0.011 | 0.002 | 5.793 | 0.155 | 0.895 |
| Morocco | Capra hircus | $2.2 \times 10^5$ | 0.011 | 0.003 | 4.192 | 0.150 | 0.627 |
| Iran | Ovis aries | $3.8 \times 10^5$ | 0.012 | 0.020 | 0.584 | 0.154 | 0.090 |
| Iran (O. orientalis) | Ovis aries | $4.5 \times 10^5$ | 0.011 | 0.021 | 0.535 | 0.148 | 0.079 |
| Iran (O. vignei) | Ovis aries | $3.7 \times 10^5$ | 0.012 | 0.020 | 0.568 | 0.151 | 0.086 |
| Various | Ovis aries | $4.1 \times 10^5$ | 0.011 | 0.021 | 0.553 | 0.152 | 0.084 |
| Morocco | Ovis aries | $4 \times 10^5$ | 0.012 | 0.022 | 0.529 | 0.153 | 0.081 |
| Barbados | Chlorocebus sabaeus | $1.1 \times 10^5$ | 0.009 | 0.002 | 5.152 | 0.170 | 0.877 |
| Central Afr. Rep. | Chlorocebus sabaeus | $1.7 \times 10^5$ | 0.009 | 0.007 | 1.266 | 0.157 | 0.199 |
| Ethiopia | Chlorocebus sabaeus | $1.4 \times 10^5$ | 0.009 | 0.002 | 3.979 | 0.161 | 0.640 |
| Gambia | Chlorocebus sabaeus | $1.4 \times 10^5$ | 0.009 | 0.008 | 1.201 | 0.159 | 0.191 |
| Kenya | Chlorocebus sabaeus | $1.5 \times 10^5$ | 0.009 | 0.003 | 2.801 | 0.165 | 0.461 |
| Nevis | Chlorocebus sabaeus | $1 \times 10^5$ | 0.009 | 0.002 | 5.361 | 0.169 | 0.908 |
| South Africa | Chlorocebus sabaeus | $1.8 \times 10^5$ | 0.009 | 0.005 | 1.912 | 0.154 | 0.294 |
| Saint Kitts | Chlorocebus sabaeus | $1.2 \times 10^5$ | 0.009 | 0.002 | 4.561 | 0.165 | 0.753 |
| Zambia | Chlorocebus sabaeus | $1.7 \times 10^5$ | 0.009 | 0.004 | 2.661 | 0.153 | 0.406 |
| African | Homo sapiens | $5.6 \times 10^4$ | 0.010 | 0.010 | 0.997 | 0.154 | 0.154 |
| Admixed American | Homo sapiens | $4.5 \times 10^4$ | 0.010 | 0.008 | 1.206 | 0.158 | 0.190 |
| East Asian | Homo sapiens | $4 \times 10^4$ | 0.010 | 0.007 | 1.455 | 0.163 | 0.237 |
| European | Homo sapiens | $4.2 \times 10^4$ | 0.010 | 0.007 | 1.316 | 0.167 | 0.219 |
| South Asian | Homo sapiens | $4.4 \times 10^4$ | 0.010 | 0.007 | 1.340 | 0.164 | 0.219 |

- $N_e$ is the estimated effective population size.
- $\mathbb{P}[\mathcal{B}_0]$ (eq. 5) is the probability for a new mutation a non-adaptive beneficial mutation. These mutations have a selection coefficient predicted at the phylogenetic-scale larger than 1, thus toward a more fit amino-acid.

- $\mathbb{P}[\mathcal{B}]$ (eq. 15) is the probability for a mutation to be beneficial. These mutations have a selection coefficient at the population-scale larger than 1.
- $\mathbb{P}[\mathcal{B} \mid \mathcal{B}_0]$ (eq. 12) is the probability for a mutation to be beneficial at the population scale, given it is predicted to be a beneficial non-adaptive mutation at the phylogenetic scale (the *precision*).
- $\mathbb{P}[\mathcal{B}_0 \mid \mathcal{B}]$ (eq. 14) is the probability for a mutation to be non-adaptive given it is beneficial at the population scale (the *recall*). This probability is obtained using Bayes' formula.

#### 11.3 Precision and recall

Table S15: Precision and recall when using a discrete DFE to estimate $S$.

| | | | **Deleterious mutations** $\mathcal{D} := S < -1$ $\mathcal{D}_0 := S_0 < -1$ | | **Nearly-neutral mutations** $\mathcal{N} := -1 < S < 1$ $\mathcal{N}_0 := -1 < S_0 < 1$ | | **Beneficial mutations** $\mathcal{B} := S > 1$ $\mathcal{B}_0 := S_0 > 1$ | |
|---|---|---|---|---|---|---|---|---|
| **Population** | **Species** | $N_e$ | **Precision** $\mathbb{P}[\mathcal{D} \mid \mathcal{D}_0]$ | **Recall** $\mathbb{P}[\mathcal{D}_0 \mid \mathcal{D}]$ | **Precision** $\mathbb{P}[\mathcal{N} \mid \mathcal{N}_0]$ | **Recall** $\mathbb{P}[\mathcal{N}_0 \mid \mathcal{N}]$ | **Precision** $\mathbb{P}[\mathcal{B} \mid \mathcal{B}_0]$ | **Recall** $\mathbb{P}[\mathcal{B}_0 \mid \mathcal{B}]$ |
| Equus c. | Equus caballus | $7.5 \times 10^4$ | 0.911 | 0.974 | 0.645 | 0.319 | 0.163 | 0.334 |
| Iran | Bos taurus | $5.6 \times 10^4$ | 0.895 | 0.973 | 0.639 | 0.288 | 0.133 | 0.283 |
| Uganda | Bos taurus | $1.3 \times 10^5$ | 0.951 | 0.974 | 0.664 | 0.492 | 0.140 | 0.159 |
| Australia | Capra hircus | $1.7 \times 10^5$ | 0.912 | 0.984 | 0.834 | 0.386 | 0.151 | 0.850 |
| France | Capra hircus | $1.9 \times 10^5$ | 0.912 | 0.983 | 0.819 | 0.381 | 0.154 | 0.940 |
| Iran (C. aegagrus) | Capra hircus | $1.9 \times 10^5$ | 0.928 | 0.980 | 0.767 | 0.408 | 0.156 | 0.911 |
| Iran | Capra hircus | $2.3 \times 10^5$ | 0.955 | 0.969 | 0.616 | 0.459 | 0.147 | 0.943 |
| Italy | Capra hircus | $1.9 \times 10^5$ | 0.911 | 0.983 | 0.822 | 0.379 | 0.155 | 0.895 |
| Morocco | Capra hircus | $2.2 \times 10^5$ | 0.950 | 0.976 | 0.706 | 0.468 | 0.150 | 0.627 |
| Iran | Ovis aries | $3.8 \times 10^5$ | 0.983 | 0.958 | 0.421 | 0.796 | 0.154 | 0.090 |
| Iran (O. orientalis) | Ovis aries | $4.5 \times 10^5$ | 0.982 | 0.961 | 0.448 | 0.806 | 0.148 | 0.079 |
| Iran (O. vignei) | Ovis aries | $3.7 \times 10^5$ | 0.974 | 0.963 | 0.480 | 0.685 | 0.151 | 0.086 |
| Various | Ovis aries | $4.1 \times 10^5$ | 0.982 | 0.964 | 0.500 | 0.822 | 0.152 | 0.084 |
| Morocco | Ovis aries | $4 \times 10^5$ | 0.983 | 0.958 | 0.387 | 0.782 | 0.153 | 0.081 |
| Barbados | Chlorocebus sabaeus | $1.1 \times 10^5$ | 0.895 | 0.987 | 0.860 | 0.351 | 0.170 | 0.877 |
| Central Afr. Rep. | Chlorocebus sabaeus | $1.7 \times 10^5$ | 0.925 | 0.974 | 0.653 | 0.373 | 0.157 | 0.199 |
| Ethiopia | Chlorocebus sabaeus | $1.4 \times 10^5$ | 0.892 | 0.980 | 0.763 | 0.320 | 0.161 | 0.640 |
| Gambia | Chlorocebus sabaeus | $1.4 \times 10^5$ | 0.939 | 0.982 | 0.770 | 0.469 | 0.159 | 0.191 |
| Kenya | Chlorocebus sabaeus | $1.5 \times 10^5$ | 0.916 | 0.976 | 0.680 | 0.345 | 0.165 | 0.461 |
| Nevis | Chlorocebus sabaeus | $1 \times 10^5$ | 0.895 | 0.989 | 0.884 | 0.358 | 0.169 | 0.908 |
| South Africa | Chlorocebus sabaeus | $1.8 \times 10^5$ | 0.930 | 0.971 | 0.604 | 0.361 | 0.154 | 0.294 |
| Saint Kitts | Chlorocebus sabaeus | $1.2 \times 10^5$ | 0.898 | 0.986 | 0.839 | 0.353 | 0.165 | 0.753 |
| Zambia | Chlorocebus sabaeus | $1.7 \times 10^5$ | 0.912 | 0.975 | 0.676 | 0.335 | 0.153 | 0.406 |
| African | Homo sapiens | $5.6 \times 10^4$ | 0.945 | 0.976 | 0.627 | 0.431 | 0.154 | 0.154 |
| Admixed American | Homo sapiens | $4.5 \times 10^4$ | 0.936 | 0.977 | 0.641 | 0.397 | 0.158 | 0.190 |
| East Asian | Homo sapiens | $4 \times 10^4$ | 0.943 | 0.977 | 0.648 | 0.422 | 0.163 | 0.237 |
| European | Homo sapiens | $4.2 \times 10^4$ | 0.945 | 0.977 | 0.644 | 0.429 | 0.167 | 0.219 |
| South Asian | Homo sapiens | $4.4 \times 10^4$ | 0.945 | 0.977 | 0.644 | 0.430 | 0.164 | 0.219 |

- $N_e$ is the estimated effective population size.
- *Precision* is the estimation of the selection coefficient at population scale ($S$) given that $S_0$ is known.
- *Recall* is the estimation of $S_0$ given selection coefficient at the population scale ($S$) is known.
- *Recall* for beneficial mutations ($\mathbb{P}[\mathcal{B}_0 \mid \mathcal{B}]$) is the probability for a mutation to be non-adaptive given it is beneficial at the population scale.

Altogether, comparing this table other estimations, we acknowledge that the exact proportion of non-adaptive among all beneficial ones is dependent on the model used to estimate the DFE. However, we can still see that non-adaptive beneficial mutations are positively selected compared to neutral and deleterious ones, and this result is not sensitive to the underlying DFE at the population scale.
